# Dorsal Hippocampus To Nucleus Accumbens Projections Drive Reinforcement Via Activation of Accumbal Dynorphin Neurons

**DOI:** 10.1101/2022.08.08.503098

**Authors:** Nicolas Massaly, Khairunisa Mohamad Ibrahim, Hye-Jean Yoon, Rossana Sandoval, Sidney Williams, Hannah Frye, William Post, Waylin Yu, Olayinka Idowu, Azra Zec, Sulan Pathiranage, Thomas L. Kash, Jose A. Morón

## Abstract

The hippocampus represents a key structure in the integration of emotional processing, learning and memory, and reward-related behaviors. While the ventral subdivision of the hippocampus (vHPC) is involved in processing emotional values of salient stimuli and goal-directed behaviors, the dorsal hippocampus (dHPC) plays a critical role in episodic, spatial, and associative memory. In addition, it has been shown that the dHPC is necessary for context- and cue-associated reward behaviors, including the expression of reward seeking. The nucleus accumbens (NAc), a central structure in the mesolimbic reward pathway, integrates the salience of aversive and rewarding stimuli and its activity is sufficient to drive aversive and appetitive behaviors. Recent evidence has demonstrated that dHPC→Nucleus Accumbens (NAc) pathway is necessary for expression of a conditioned place preference. However, despite years of groundbreaking research and identification of direct projections from the dHPC to the NAc, the sufficiency for dHPC→NAC inputs to drive reinforcement and reward associated behavior remains to be determined.

Here using a wide range of complementary and cutting-edge techniques including behavior, *in-vivo* manipulation using optogenetics, chemogenetics, brain clearing, local field potential and fiber photometry recordings, we demonstrate that activation of excitatory projections from the CA1 subregion of the dHPC (dCA1) is sufficient to drive reinforcing behaviors. In addition, we provide strong evidence that this reinforcing behavior is driven by 1) a direct projection from the dCA1 to the NAc and 2) enhanced glutamatergic signaling within the NAc. Furthermore, we uncovered that while dCA1 stimulation increases the activity of both enkephalin- and dynorphin-containing medium spiny neurons in the NAc, the selective activity of dynorphin-containing neurons is necessary for the expression of this reinforcing behavior. Our findings shed light on a novel pathway governing reinforcement and further extend the role of the dHPC on reward seeking.

## INTRODUCTION

Despite years of groundbreaking research, the exact circuits and brain structures involved in governing reinforcement and goal-directed behaviors are still not fully uncovered. The hippocampus is an heterogenous brain structure, in which the dorsal subdivision (dHPC) is critically involved in spatial and contextual memory while its ventral subdivision (vHPC) integrates emotional value of salient stimuli and guides goal directed behavior ^1–5^. Indeed, vHPC activity modulates both rewarding stimuli, drug relapse and threat avoidance behaviors ^6–13^, confirming its role as a key structure in emotional encoding ^2,14,15^. On the other hand, the dHPC plays a crucial role in the integration, maintenance, and retrieval of contextual, spatial and reward-associated memories ^16–25^. We and others have demonstrated that dHPC activity is necessary to initiate goal-directed behavior to avoid stressor/threat or engage in reinforcement in response to cues/contexts previously associated with an aversive or rewarding stimulus, respectively ^1,1,19,20,25–29^. These findings highlight a critical role for the dHPC in reward processing.

The nucleus accumbens (NAc), a central structure of the mesolimbic dopaminergic pathway, plays a critical role in integrating dopaminergic reinforcement signals from the ventral tegmental area (VTA) ^30–32^. Previous investigations have established that excitatory CaMKII+ afferents to the NAc including inputs from the amygdala, the prefrontal cortex and the vHPC are sufficient to drive reinforcement ^33^. While the vHPC is sufficient to reinforce instrumental behavior, more recent findings have demonstrated that CaMKII+ afferent from the CA1 region of the dHPC (dCA1) to the NAc Shell (NacSh) are necessary for the expression of contextual memories induced by salient stimuli ^23^. In addition, early evidence uncovered that dHPC electrical stimulation is sufficient to elicit reinforcement independently of associative learning ^34,35^. However, the current literature still lacks clear evidence assessing the sufficiency for direct dCA1 → NAc projection to drive reinforcement, a critical component of reward seeking and expression of reward-associated memories.

Here, we used a combination of optogenetic and chemogenetic approaches in freely moving animals, *in-vivo* local field potentials and fiber photometry to investigate the sufficiency of dCA1 excitatory projection neurons to the NAcSh to drive reinforcement via local glutamatergic signaling acting at cell-specific neuronal types. Using an optogenetic self-stimulation instrumental procedure, we demonstrate that recruitment of dCA1 CaMKII+ projection neurons to the NAcSh is sufficient to drive instrumental reinforcement. Moreover, we uncover that dCA1 stimulation is associated with a significant increase in calcium transient, a proxy for neuronal activity, in both dynorphin- and enkephalin-containing medium spiny neurons (MSNs) within the NAcSh. Lastly, we demonstrate that dynorphin-containing MSNs are necessary to drive dCA1 CaMKII+-induced reinforcement. Altogether, our data suggest that the dCA1 pyramidal projection neurons to the NAcSh are sufficient to drive reinforcement and trigger reward seeking via dynorphin-containing MSNs activity.

## RESULTS

### Stimulation of excitatory neurons in the dCA1 drives reinforcement

To investigate the sufficiency for dCA1 excitatory neurons in driving reinforcing behaviors we used an operant light self-stimulation procedure (**Figure 1A**). In order to target dCA1 excitatory neurons, mice were injected with viral construct carrying ChR2 under CaMKII promotor (AAV5-CaMKII-ChR2-eYFP). Two weeks after viral infusion, mice were permanently implanted with a fiber optic right above the dCA1. A week after surgery, mice were placed in an operant chamber, and a light stimulation (20 Hz, 15 ms pulses, 4 mW for 2 s) was delivered with every nose poke in an active (illuminated) inset (Fixed Ratio 1; FR1) followed by a 10 second time out. On the first training day, animals expressing ChR2 in dCA1 CaMKII+ neurons significantly sought light self-stimulation. This behavior was maintained throughout the whole procedure, as demonstrated by the significant difference in the number of nose pokes into the active insets versus the inactive insets across days (**Figure 1B**). On the contrary, animals injected with a control virus (control group) did not discriminate between the active and inactive insets, confirming the reinforcing nature of dCA1 CaMKII+ neuronal activation (**Figure 1B**). After stable self-stimulation behavior observed across days (**Figure 1B)**, the same cohort of mice underwent extinction sessions during which all cues were presented, but interaction with either nose insets had no further consequence (laser turned OFF). During extinction sessions, mice expressing ChR2 in dCA1 CaMKII+ neurons significantly decreased their interaction with the active insets, uncovering the necessity for CaMKII+ neurons activation to maintain reinforcement (**Figure 1C**). Lastly, 24 hours after the last extinction session, mice were exposed to a “reinstatement session” during which the laser was turned back ON and a nose poke in the active inset triggered light-stimulation of dCA1 CaMKII+ neurons. During this test, ChR2 expressing animals demonstrated a significantly higher interaction with the active inset than the inactive inset (**Figure 1D**). In addition, the ChR2 expressing animals also exhibited a significantly higher number of interactions with the active inset as compared to control littermates (**Figure 1D**). Overall, these findings uncover the sufficiency for dCA1 CaMKII+ activation to trigger light self-stimulation.

**Figure 1:**
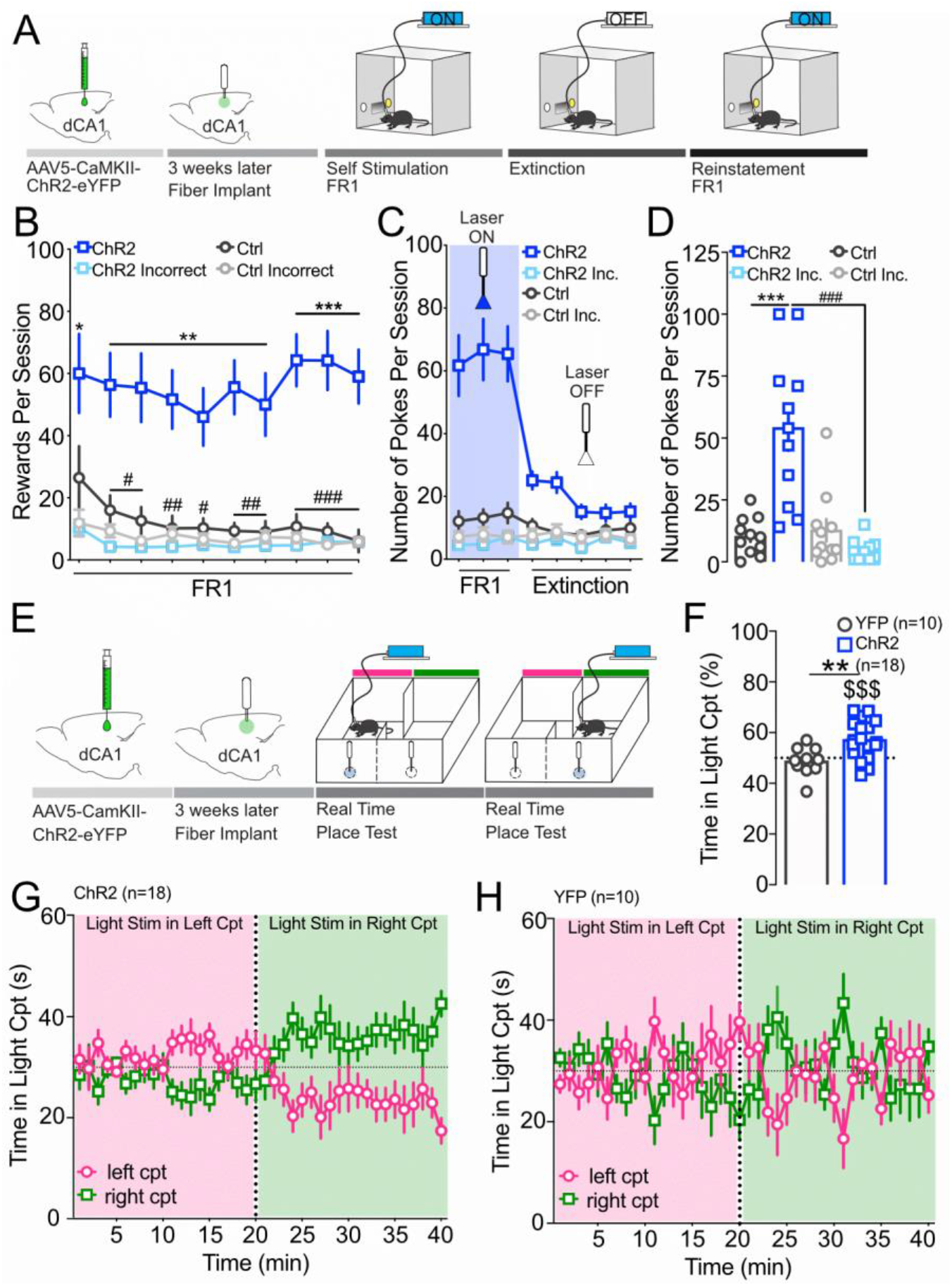
Stimulation of CaMKII+ neurons in the dCA1 drives reinforcement. A) Representative schematic of the behavior. B) Animals expressing ChR2 within the dCA1 (n=11) rapidly discriminated between the active and inactive nose insets, seeking light-stimulation through persistent active nose pokes. C) Mice stopped seeking for light stimulation when an active nose-poke was not associated with light delivery within the dCA1 (Extinction). D) 24 hours after the last extinction session, ChR2 expressing mice (n=11) demonstrate a significantly higher number of active nose pokes than control littermates (n=11) (p < 0.001) and their incorrect nose pokes (p < 0.001), suggesting that active seeking is dependent on dCA1 CaMKII+ neurons stimulation. E) Representative schematic of the behavior. Briefly, a week after viral expression and securing a fiber implant above the dCA1 animals were exposed to an RTPT procedure in which light-associated compartment changed at the 20 minutes mark of the unique 40 minutes test. F) Animals expressing ChR2 in CaMKII+ in the dCA1 (n=18) exhibit a significant overall preference for the light associated compartment. G) Animals expressing ChR2 in the dCA1 show a preference for the left compartment within the first 20minutes of the test (pink) when light stimulation is associated with animal’s presence in the left compartment. Mice’s preference changes towards the right compartment during the last 20 minutes of the test (green) when light stimulation is associated with animal’s presence in the right compartment. H) In sharp contrast, control animals injected with eYFP control virus (n=10) within the dCA1 do not exhibit any preference for any compartment within the 40 minutes session. Data are expressed as mean +/-S.E.M. * p<0.05, ** p < 0.01 and *** p < 0.001 Two-way ANOVA followed by post hoc Tukey comparing control versus ChR2 animals, ### p<0.001 post hoc Tukey comparing active versus inactive nose-pokes, and $$$$ p < 0.0001 One Sample T-test comparing percent in stimulation chamber to 50% as a hypothetical value.

Interestingly, the sole presentation of the cue associated with light self-stimulation during the 10 first days of self-stimulation was not sufficient to trigger reinforcement (extinction period), confirming that CaMKII+ activation is a necessary driver for seeking behavior. To further confirm the reinforcing properties of CaMKII+ neurons stimulation, we used an unbiased custom-made real-time place test (RTPT) assay composed of 2 black matte compartments with no external cues or context (**Figure 1E-H**). In this procedure, the animal’s presence in one compartment of the apparatus triggered light delivery while the presence in the other had no further consequence. In addition, to ensure that the preference for light-paired compartment was not simply due to alterations in exploratory behavior but rather active seeking for light stimulation, the light-paired and neutral compartments were reversed within the session (at the 20 minutes mark of the 40 minutes test – **Figure 1E**). ChR2-expressing mice actively sought out light-stimulation when the light-paired compartment was changed, suggesting that light-stimulation preference was due to active seeking for CaMKII+ neurons stimulation rather than decreased exploratory behavior (**Figure 1F and G**). In sharp contrast, control littermates did not develop any preference for either compartment across the 40-minute test (**Figure 1F and H**). Overall, our experiments thus far demonstrate that activation of CaMKII+ neurons within the dHPC is sufficient to reinforce instrumental behavior.

### dCA1 CaMKII+ neurons are anatomically and functionally connected with the NAc

Recent evidence has uncovered that CaMKII+ dCA1 projections onto the NAcSh are necessary for the expression of contextual memories induced by appetitive stimuli ^23^. Here, we used a brain clearing approach following AAV1.Syn.NES.jRGECO1a.WPRE.SV40 injection within the dCA1 to determine downstream projections of CaMKII+ neurons (**Figure 2A, B**). In agreement with previous reports, we demonstrated that the dCA1 sends direct projections to the mesolimbic pathway, including the NAc and the VTA, amongst other downstream structures (**Figure 2B**). In a separate cohort of mice, AAV5-ChR2-CaMKII-eYFP was expressed in the dCA1, and the coronal slices showed that dCA1 CaMKII+ neuronal projections mostly target the most rostral part of the NAc as previously reported (**Figure 2C**) ^23^. However, recordings assessing the connectivity between these two brain areas in freely moving animals has not been provided yet. To assess the *in-vivo* connectivity between dCA1 CaMKII+ projecting neurons and the NAc, we coupled an optogenetic approach together with local field potential (LFP) recordings. Briefly, after injecting CaMKII+ promoter driven ChR2 in the dCA1 of mice, we implanted a custom-made 3D printed headcap that held a fiber implant above the dCA1 region of the hippocampus and 16 electrodes (8 electrodes on each side) within the rostral part of the NAc. Mice were then placed in an open field area and LFP were recorded in response to dCA1 CaMKII+ neurons stimulation (5 ms duration, every 5 s, 4 mW) (**Figure 2D**). Overall, 87.5 % of the electrodes implanted in the NAc recorded evoked LFP with dCA1 stimulation (**Figure 2E**). Furthermore, a significant increase in overall LFP amplitude recorded was observed upon stimulation of dCA1 CaMKII+ neurons (**Figure 2F**). The evoked activation of the NAc under dCA1 stimulation provides further evidence of a functional dCA1-NAc circuitry that may underlie the reinforcing properties of dCA1 CaMKII+ neurons. To further determine which subdivision of the rostral NAc was connected to dCA1 stimulation and gain both anatomical and temporal resolution of the dynamics of NAc neuronal activity following stimulation of the dCA1, we used *in-vivo* fiber photometry and recorded NAcSh calcium transients as a proxy for neuronal activity. We expressed a calcium indicator under an ubiquitous synapsin promoter within the NAcSh (AAV-hSyn-GCamp6f-eYFP) and ChR2 in dHPC CaMKII+ neurons; a fiber optic was placed above the dCA1 to locally stimulate ChR2-expressing neurons (**Figure 2G**). Under these conditions, we demonstrated that dCA1 CaMKII+ neuron self-stimulation is sufficient to evoke a significant increase in calcium transients within the NAcSh (**Figure 2H-J**). No differences in calcium transients were observed with light stimulation in mice that expressed the control virus in the NAcSh (**Figure 2H-J**), and no change in calcium transients were observed in response to the presentation of the cue light during the self-stimulation procedure (**Suppl. Fig 1A-D**). Interestingly, similar results were obtained when CaMKII+ recruitment was triggered by the experimenter (non-contingent light delivery) (**Suppl. Fig 1E-H**), confirming that NAc Ca2+ transients are indeed triggered by dHPC CaMKII+ neurons activity. These results further reveal a selective accumbal response to dCA1 CaMKII+ neuron stimulation.

**Figure 2:**
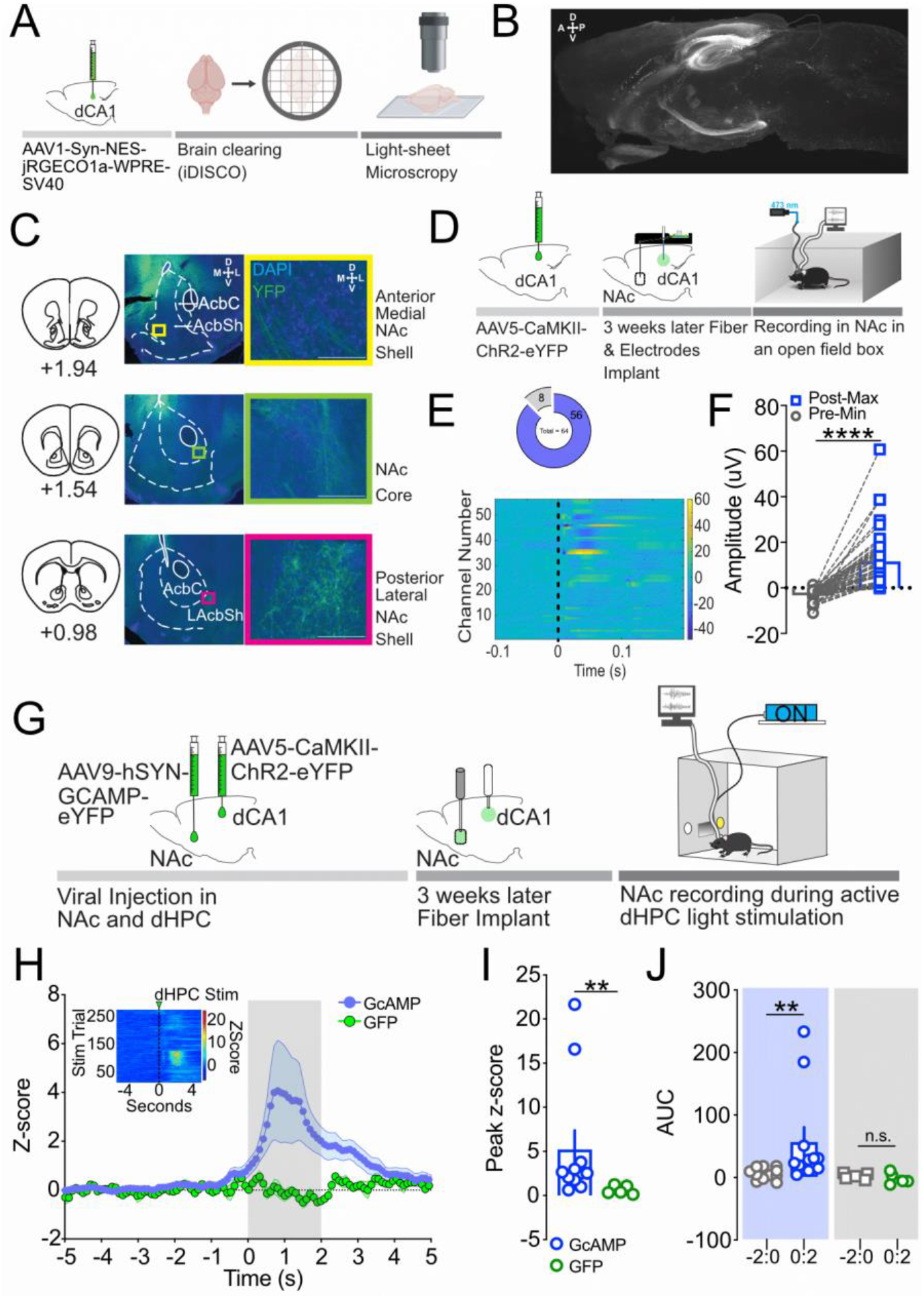
dCA1 CaMKII+ neurons stimulation is functionally connected with the NAc. A) Schematic representation of the brain clearing procedure. B) Sagittal section of a mice brain representing dCA1 projections throughout the brain. C) In a separate cohort, coronal sections of brains injected with CaMKII promoter driven ChR2 in the dCA1 were used to determine anatomical projections to the NAc. D) Schematic representation of the *in-vivo* local field potential procedure. E) Functional connectivity with 56 electrodes in the NAc out of 64 showed responsiveness to dHPC stimulation (upper panel). Heat map of the recordings obtained for a 0.3 s window around dCA1 CaMKII+ neurons stimulation (t = 0) (lower panel). F) A significant increase in absolute evoked LFP amplitude in the 56 electrodes implanted in the NAc that responded with stimulation of dCA1 CaMKII+ neurons. G) Schematic representation of the self-stimulation procedure. H) Time course for the Z-scores of the calcium transient. Gray area represents dCA1 self-stimulation. Calcium transients selectively increase in animals self-stimulating and expressing both ChR2 in the dCA1 and GCaMP6f in the NAcSh. Insert: heatmap of the calcium transient upon light self-stimulation. I) Peak Z-score during the 2 seconds of dCA1 CaMKII stimulation. Peak Z-score is significantly higher in ChR2-GCaMP6f expressing animals. J) Area under the curve (AUC) of 2 seconds prior and during dCA1 light stimulation obtained in experimental groups. The measured calcium transients AUC in the NAcSh of ChR2-GCaMP6f animals upon light self-stimulation (ChR2-GCaMP: 0:2) were significantly higher than their respective baseline (ChR2-GCaMP: -2:0). The calcium transients AUC in the control group not expressing GCaMP in the NAcSh (ChR2-GFP) did not significantly change upon light stimulation. All data are expressed as mean +/-S.E.M. and n.s. p > 0.05; ** p< 0.01; **** p < 0.0001. Paired T-test comparing the min and max of evoked LFP amplitude, Mann-Whitney test for peak z-score between GcAMP and GFP groups and, Wilcoxon matched-paired rank tests for AUC in 0:2 and its respective baseline -2:0.

### dCA1 to NAcSh projections are sufficient to drive reinforcement via glutamatergic transmission

While stimulation of dCA1 CaMKII+ neurons is sufficient to trigger both a LFP response and an increase in calcium transients in the NAcSh, the sufficiency for activation of dHPC terminals in the NAcSh to drive reinforcement remained to be determined. To assess this, we expressed ChR2 in CaMKII+ neurons within the dCA1 and placed a fiber implant above the NAcSh (**Figure 3A**). A week after securely positioning the fiber implant and to allow full recovery, animals were exposed to a Fixed Ratio 1 (FR1) schedule of reinforcement during which an interaction with the active nose inset triggered light-stimulation in the NAcSh (**Figure 3A**). Similar to the results obtained with somatic activation of CaMKII+ neurons in the dCA1, ChR2 expressing animals rapidly discriminated between the active and inactive port of the self-administration apparatus (**Figure 3A**). During extinction sessions, where the lasers were turned off, the same cohort of mice significantly decreased their interaction with the active insets, uncovering the necessity for dHPC CaMKII+ terminal stimulation in the NAcSh to maintain reinforcement (**Figure 3B**). Furthermore, when exposed to a “reinstatement session” during which the laser was turned back ON, ChR2 expressing animals had a significant increase in their interaction with the active inset as compared to both the inactive inset and the active inset during the last extinction session (**Figure 3C**). The obtained results uncovered that stimulation of dHPC terminals within the NAc is sufficient drives reinforcing behavior.

**Figure 3:**
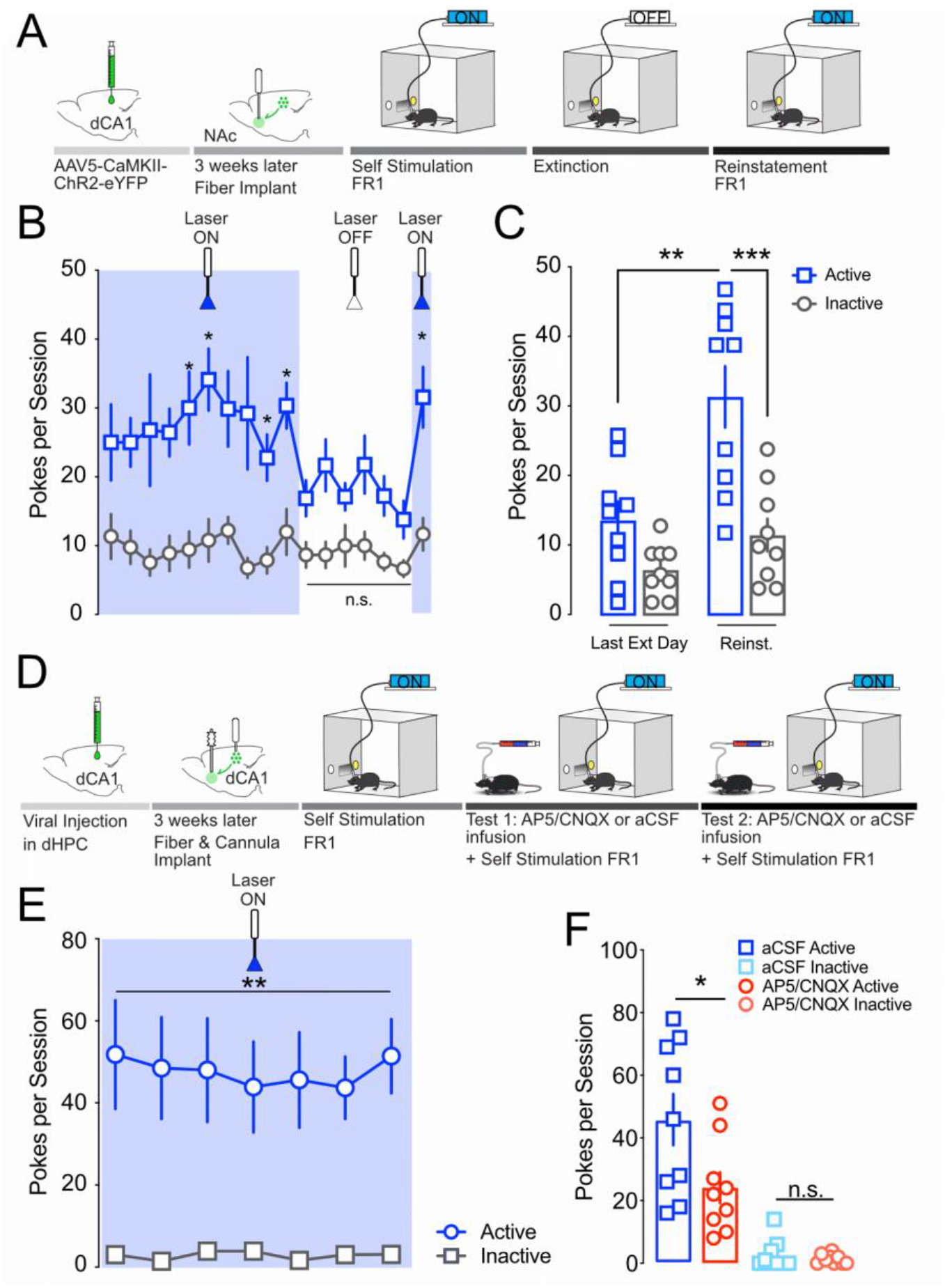
Projections from the dCA1 selectively to the NAcSh are sufficient to drive reinforcement. A) Schematic representation of the surgeries and self-stimulation procedure. B) Mice expressing ChR2 within the dCA1 but received light stimulation in the NAcSh (n=9) rapidly discriminated between the active and inactive insets, seeking light stimulation through persistent active nose pokes. The persistence of reinforcing behavior stopped when the active nose-poke was not associated with light delivery (extinction sessions; laser turned OFF). C) 24 hours after the last extinction session, animals were exposed to a reinstatement session during which the laser was turned back ON. Mice interacted significantly more with the active than the inactive inset (p <0.001). In addition, there was significantly higher interaction with the active inset during the reinstatement session than the last extinction session (p <0.01). D) Schematic representation of the surgeries and self-stimulation procedure. E) During training animals (n=10) rapidly discriminated between the active and inactive insets and maintained stable self-stimulation across days. F) Local micro-infusion of AP5/CNQX cocktail within the NAcSh to block glutamatergic signaling significantly decreased active inset interactions as compared to aCSF micro-infusion. Data are expressed as mean +/-S.E.M. Two-way ANOVA for repeated measures followed by Sidak’s multiple comparison post-hoc tests, n.s. p > 0.05; ** p < 0.001; *** p < 0.001; and **** p< 0.0001.

To further investigate whether the reinforcing properties of dHPC→NAcSh activation are mediated by local glutamatergic transmission within the NAcSh, we combined opto-stimulation of CaMKII+ hippocampal neurons with local pharmacology. After infusion of CaMKII+ driven ChR2 in the dCA1, implantation of fiber implant above the dCA1 and guide-cannula above the NAcSh, animals were trained to self-stimulate (**Figure 3 D**). Once the reinforcing behavior was stable (**Figure 3E**), mice were infused with either a cocktail of AMPA/NMDA antagonists (CNQX:0.7mM/AP5:1.6mM) or aCSF as a control in the NAcSh 30 minutes prior to an FR1 self-stimulation session ^36^. Each animal received both treatments 3 days apart in a counter-balanced manner (half received CNQX/AP5 before the first test, while the other half received aCSF). Blockade of glutamatergic signaling within the NAcSh blunted self-stimulation, as shown by a significant decrease in interactions with the active inset after CNQX/AP5 infusion as compared to aCSF infusion (**Figure 3F**). Altogether our data demonstrate that dCA1→NAcSh CaMKII+ projections are sufficient to drive reinforcement through local glutamatergic signaling in the NAcSh.

### Accumbal dynorphin containing neurons are necessary for dCA1 driven reinforcement

The NAc is composed of 90 ∼ 95 % GABAergic MSNs organized in two distinct peptidergic populations, enkephalin- and dynorphin-containing MSNs, which have been identified and studied for their role on rewarding and aversive behaviors ^37,38^. We used transgenic mice models to assess whether CaMKII+ neurons in the dHPC are preferentially recruiting one MSN population over the other to drive reinforcement. Enkephalin-IRES-Cre or Dynorphin-IRES-Cre mice were injected with a Cre-dependent calcium indicator (AAV-Syn-Flex-GcAMP6f-eYFP) in the NAcSh and ChR2-CaMKII (AAV5-CaMKII-ChR2-eYFP) in the dCA1 (**Figure 4A**). Both enkephalin- and dynorphin-containing neurons in the NAcSh exhibited a significant increase in calcium transients upon light stimulation of the dCA1 CaMKII+ neurons as compared to their respective baseline. No significant differences were observed between the two cell populations (**Figure 4B-D**). To dissect the necessity of NAcSh cell types in dCA1-mediated reinforcement we used a chemogenetic inhibitory approach. Enkephalin-cre and dynorphin-cre mice were injected with a Gi DREADD (AAV5-hSyn-hM4Di-mCherry or AAV5-hSyn-mCherry as control) in the NAcSh and CNO (1mg.kg^-1^, i.p.) was injected 30 minutes before the self-stimulation sessions (**Figure 4E**). Silencing the activity of dynorphin-containing neurons during the light self-stimulation sessions is sufficient to significantly reduce reinforcement (i.e., number of nose pokes in the active inset) (**Figure 4F, H, Suppl Figure 2C**). In sharp contrast, silencing enkephalin containing neurons did not impact self-stimulation behavior (**Figure 4G-H, Suppl Figure 2B**). Altogether, we uncovered that while dCA1 stimulation increases the activity of both enkephalin and dynorphin MSNs within the NAcSh, the selective activity of dynorphin-containing neurons is necessary for the expression of reinforcing behavior.

**Figure 4:**
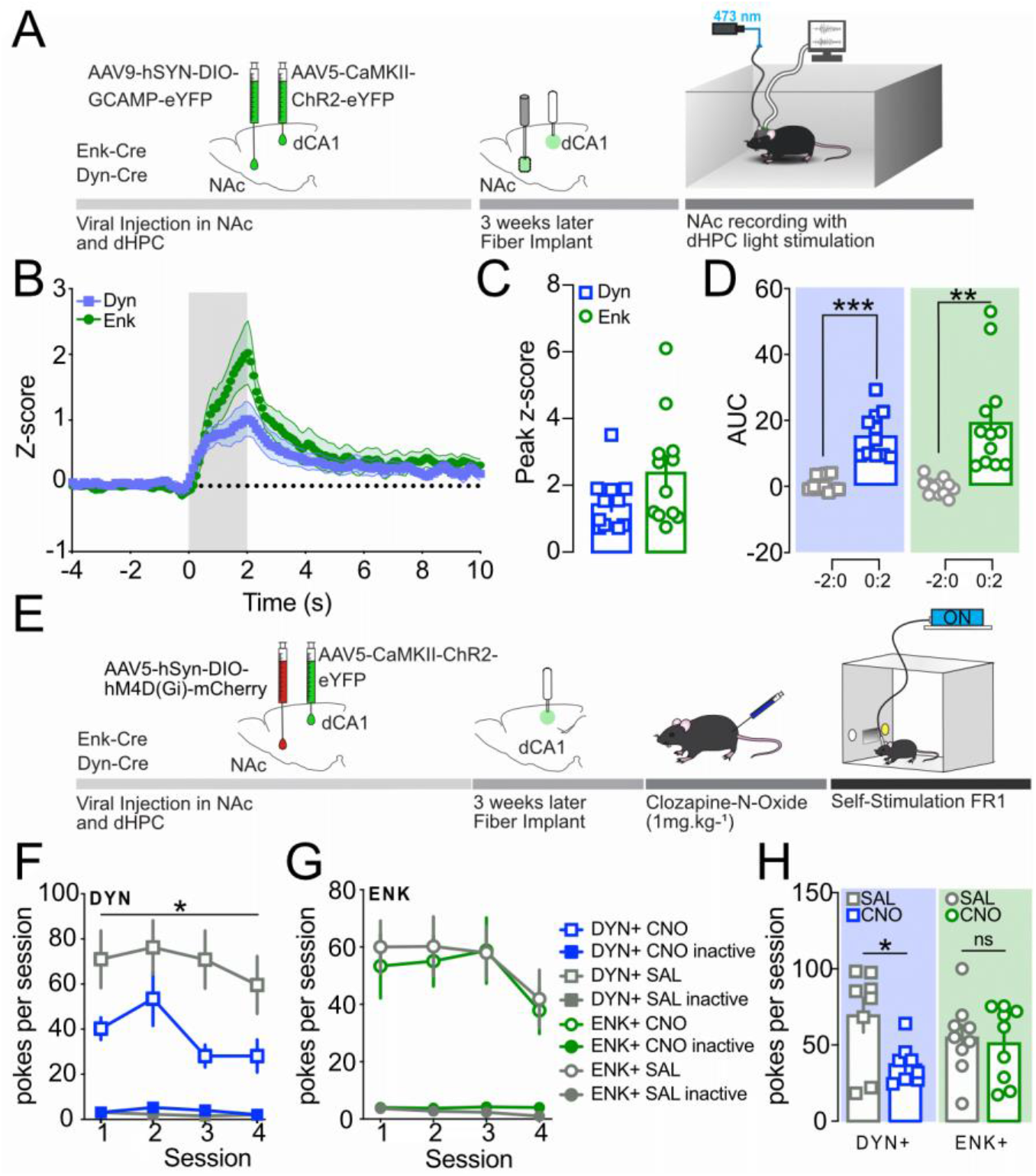
Accumbal dynorphin containing neurons are necessary for dCA1 CaMKII+ neurons driven reinforcement. A) Schematic representation of the performed surgeries and non-contingent (experimenter-induced) stimulation procedure. B) Time course for the Z-scores of the evoked calcium transient. Gray area represents dCA1 stimulation. Calcium transients selectively increase in both enkephalin- and dynorphin-cre animals upon stimulation of CaMKII+ neurons. C) Peak Z-score during the 2 seconds of dCA1 CaMKII+ neurons stimulation. Peak Z-score does not significantly differ between dynorphin (n=11) and enkephalin (n=12) neuronal recordings upon dCA1 CaMKII+ neurons stimulation. D) Area under the Curve (AUC) for z-scores obtained in experimental groups demonstrate that both enkephalin and dynorphin calcium transients significantly increase during stimulation (0:2) compared to their respective baseline (−2:0). E) Schematic representation of viral injection, surgical procedure, and behavioral protocol. F) All animals have been exposed to 4 saline pretreatment and 4 CNO pretreatment sessions. Silencing dynorphin containing neurons within the NAcSh significantly reduces light self-stimulation (n=8). G) All animals have been exposed to 4 saline pretreatment and 4 CNO pretreatment sessions. Silencing enkephalin containing neurons within the NAcSh does not impact light self-stimulation (n=9). H) Overall number of self-stimulations is significantly reduced selectively when dynorphin-containing neurons are silenced within the NAcSh, uncovering the necessity of those neurons to drive dCA1 CaMKII+-induced reinforcement. Data are expressed as mean +/- S.E.M. and n.s. p > 0.05; * p < 0.05; ** p<0.01; *** p < 0.001. Unpaired Mann-Whitney test for peak z-score between Dyn and Enk groups; paired T-test for AUC in 0:2 and its respective baseline -2:0, Two-way ANNOVA RM comparing effects Saline vs CNO pretreatment in Dyn and Enk groups, respectively and, paired T-test comparing active pokes with Saline and CON pretreatment in both Dyn and Enk mice, respectively.

## DISCUSSION

In the current study, we used light self-stimulation to demonstrate that stimulation of CaMKII+ neurons within the dCA1 is sufficient to drive reward seeking behavior and reinforcement. This finding uncover cell specificity to mediate dHPC-induced reinforcing behaviors identified by early studies using electrical stimulation ^34,35^. Despite many reports demonstrating the role of hippocampal pyramidal cells in encoding location, context, cues and memory expression associated with appetitive and aversive behaviors ^1,22,23,39–41^, no study to date has assessed the sufficiency for dCA1 CaMKII+ neurons to drive reinforcement.

Appetitive rewards such as sucrose can trigger dHPC activity and engage the NAc to encode guided appetitive behaviors ^23,42–45^. However, these previous studies lacked *in-vivo* measures of dHPC to NAc connectivity and did not provide any insights into the potential role for excitatory transmission from dHPC terminals onto NAc synapses to drive reinforcing behavior. Our data demonstrates that self-stimulation of dCA1 CaMKII+ neurons is reinforcing and leads to significant increases in calcium transients in the NAcSh (**Figure 2G-J, Suppl Figure 1E-H**). Thus, our results further expand on the critical role for dorsal hippocampal-accumbens functional connectivity in driving reward encoding ^23,44,45^.

The NAc receives dense excitatory projections including dopaminergic inputs from the VTA and glutamatergic excitatory inputs from cortical and subcortical structures, such as the amygdala, the prefrontal cortex and the vHPC ^33,46–48^. Recently, Trouche et al (2019) uncovered that a direct monosynaptic circuit between dCA1 CaMKII+ neurons and NAc paravalbumin interneurons and MSNs neuronal populations is necessary for the expression of appetitive memory ^23^.

Additionally, they demonstrate that silencing those projections do not impact reward foraging in their CPP assay, confirming that silencing dCA1→NAC impairs the expression of conditioned place preference rather than reward seeking itself. On the other hand, our study demonstrates that activation of dCA1→NAc is sufficient to promote reward seeking, providing an additional level of complexity to the role of this pathway.

We used *in-vivo* fiber photometry to determine the extent to which dCA1 recruitment could impact accumbal activity. We demonstrate that stimulation of dCA1 CaMKII+ neurons, either during “reward-seeking” behavior in a contingent light self-stimulation procedure or when light is triggered non-contingently by the experimenter, is sufficient to significantly increase accumbal activity, suggesting an engagement of the NAcSh in dCA1-induced reinforcement. Previous evidence has shown that CaMKII+ projection neurons from the dCA1to the NAc engage paravalbumin fast spiking interneurons (PV+FSI) and, to a lower extent, MSNs ^23^. In this context, the activity of the latter is tightly controlled by PV+FSI directly recruited by dCA1 afferent. NAc MSNs are typically classified in two main population according to their opioid peptidergic content (dynorphin versus enkephalin) and dopamine receptor expression (D1 versus D2 receptor); dynorphin containing MSNs vastly expressing D1 receptors while enkephalin-containing MSNs express mostly D2 receptors ^37,48,49^. Several reports have demonstrated that the activity of dynorphin-containing (D1R) and enkephalin-containing (D2R) MSNs play a critical role in rewarding or aversive experience, respectively ^37,38,50–56^. However, these results are in conflict with other reports demonstrating the recruitment of D2 (enkephalin) MSNs during appetitive/rewarding tasks ^44,57^ and D1(dynorphin) MSNs necessity to drive aversive behavior such as negative affect and anhedonia ^11,38^. Thus, considering the dichotomic roles of those two neuronal populations within the NAc, we pursued a cell-specific approach to further investigate the necessity of each MSNs subtype on dHPC CaMKII+ driven reinforcement.

Our findings demonstrate that CaMKII+ neuron stimulation within the dCA1 triggers an increase in calcium transients in both enkephalin- and dynorphin-containing MSNs within the rostral part of the NAcSh. While these two distinct neuronal populations responded similarly to dCA1 light-stimulation, only selective silencing of dynorphin-containing MSNs was sufficient to dampen dCA1-induced reinforcing properties as indicated by a reduction in self-stimulation of dCA1 CaMKII+ neurons. In contrast, silencing enkephalin-containing MSNs had no effect on CaMKII+-driven reinforcement. Interestingly, Al-Hasani et al. demonstrated that activity of dynorphin-containing MSNs can both trigger appetitive or aversive behavioral outcome depending on the ventro-dorsal anatomical distribution of the recruited neurons ^49^. Similarly, Castro and Berridge have uncovered a rostro-caudal axis defining hot and cold spots areas within the NAcSh for which local infusion of the same opioid agonists can drive either an increase or decrease in “liking” behaviors in rats ^58^. Further investigation on the dichotomic effects of MSNs in the NAc is needed in order to 1) properly assess the role of those enkephalinergic and dynorphinergic populations along the dorso-ventral and rostro-caudal axes within the NAcSh and 2) reconcile current conflicting results on the rewarding and aversive roles of NAcSh MSNs. In addition, the involvement of PV+FSI and local MSNs collaterals in finely tuning the activity of each MSN subtype underlying the dCA1→NAcSh reinforcement remains to be fully elucidated.

Overall, our study brings an additional level of granularity to the role of dHPC in reward seeking and memory processes. While previous studies have described the direct pyramidal dCA1→NAc projections ^23,45,59^ and their role in expression of contextual memories ^23^, we demonstrate here that activating this pathway is also sufficient to drive reinforcement via the activity of dynorphin-containing neurons within the NAcSh. We believe that the reinforcing properties of dCA1→NAc projections, in addition to their role in cue and contextual memory, play a critical part in the expression of cue- and context-dependent reward seeking.

Overall, these findings have a very high potential for significant impact in the field, as we uncovered that hippocampal activity governs reinforcing behaviors via direct projections to the NAcSh and via the activity of dynorphin-containing neurons. By further characterizing the specific brain circuits involved in this behavioral adaptation, this study highlights a role for the dHPC→NAc pathway in the expression of reward seeking including relapse and drug use disorders.

## METHODS

### CONTACT FOR REAGENT AND RESOURCE SHARING

Further information and requests for resources and reagents should be directed to and will be fulfilled by the Lead Contact, Jose A. Morón.

### EXPERIMENTAL MODEL AND SUBJECT DETAILS

All procedures were approved by the Washington University Institutional Animal Care and Use Committee (IACUC) in accordance with the National Institutes of Health Guidelines for the Care and Use of Laboratory Animals. Adult WT, PEnk-IRES-Cre, and PDyn-IRES-Cre C57BL/6J male and female mice (20-30 g) were used for this study. All animals were 10 to 12 weeks at the beginning of the experiments. Four to five mice were housed together, given access to food pellets and water ad libitum, and maintained on a 12/12 h dark/light cycle (lights on at 7:00 AM). All animals were kept in a sound-attenuated, isolated holding facility in the lab 1 week prior to surgery, post-surgery, and throughout the duration of the behavioral assays to minimize stress.

### SURGERIES

All surgeries were performed under isoflurane (1.5/2 MAC) anesthesia under sterile conditions.

Mice were anesthetized in an induction chamber (4 MAC Isoflurane) and placed into a stereotaxic frame (Kopf Instruments, Model 1900) where they were maintained at 1.5 MAC–2 MAC isoflurane. A craniotomy was performed and followed by a unilaterally injection, using a blunt needle (86200, Hamilton Company), 500 nl of AAV5-CaMKII-ChR2-eYFP or, AAV5-CaMKII-eYFP (Hope Center Viral Vector Core, viral titer 2.8 × 10^13^ vg/mL) into the dHPC (stereotaxic coordinates from Bregma: A/P: – 1.70 mm, M/L: ± 1.5 mm, D/V: –1.80 mm) ^60^. For chemogenetic approaches, animals were injected with 250 nl of AAV5-hSyn-hM4Di-mCherry (Addgene, viral titer 2.8 × 10^13^ vg/mL) in the NAcSh (stereotaxic coordinates from Bregma: A/P: + 1.70 mm, M/L: ± 0.5 mm, D/V: –4.50 mm) during the same procedure. For fiber photometry experiments, a calcium indicator AAV9-hSyn-GCAmp6f-eYFP (Addgene, viral titer 2.8 × 10^13^ vg/mL) or a control virus AAV9-hSyn-eYFP (Addgene, viral titer 2.8 × 10^13^ vg/mL) was injected in the NAcSh of WT mice while AAV9-Syn-Flex-GcAMP6f-eYFP (Addgene, viral titer 2.1 × 10^13^ vg/mL) was injected in the NAcSh of cre-dependent mice (stereotaxic coordinates from Bregma: A/P: + 1.70 mm, M/L: ± 0.5 mm, D/V: – 4.50 mm). Three weeks after intracranial local injections, mice underwent a second surgery during which a fiber optic was implanted in the dHPC, and also above the NAcSh for fiber photometry experiments. Alternatively, a bilateral guide cannula was implanted 0.5mm above the NAcSh (AP: +1.7mm; ML: +/-0.5mm; DV: -4.25mm) to allow local AMPA/NMDA antagonists infusion. The implants were secured with Metabond dental cement using Radiopaque L-powder, B Quick Base, and C Universal TBB Catalyst (Parkell). Mice were allowed to recover for at least 1 week before running any behavioral experiment.

Furthermore, the experiments also started 4 weeks after the adeno-associated virus injection, permitting optimal expression of ChR2 in the infected neurons.

### CHEMOGENETICS, OPTOGENETICS AND BEHAVIORAL ASSAYS

#### LIGHT SELF-STIMULATION

Mice operant-conditioning chambers (Med Associates, Fairfax, VT) were equipped with nose insets accessible to animals. Mice were exposed to a Fixed Ratio (FR) session in which fiber implant was connected to a 473 nm laser. A cue light was presented on the active inset and active interaction with this port resulted in a laser activation (20 Hz, 15 ms pulses for 2 s) together with cue light being turned off for 10 s (time out period) and any interactions with the insets during this period had no consequences. Interaction with the inactive inset also had no consequences. Mice were trained to discriminate between active and inactive insets during the 1-hour long FR1 sessions. In **Figure 1** and **3A-C**, animals underwent extinction sessions for 5 days during which all parameters where kept constant but interaction with the active inset did not result in light stimulation (laser was turned OFF). After animals extinguished their discrimination between the active and inactive insets, they underwent a single reinstatement procedure (FR1) during which interaction with the active inset led to the delivery of light stimulation.

In **Figure 3D-F**, the animals were first trained to self-stimulate before receiving intra-NAcSh infusion of either a cocktail of AMPA/NMDA antagonists (CNQX:0.7mM/AP5:1.6mM) or aCSF (control) 30 minutes prior to an FR1 self-stimulation session ^36^. The treatments were counterbalanced, where half of the animals received CNQX/AP5 30 minutes before the first test while the other half received aCSF. The second test was done 3 days after the first test where the local infusion of drugs was switch between the groups (i.e previously received aCSF mice in first test will receive CNQX/AP5 in the second test).

In **Figure 4**, after training sessions all animals were exposed to 8 days of 1-hour self-stimulation sessions. For 4 out of the 8 consecutive days mice received an intraperitoneal injection of CNO (1mg.kg^-1^) 30 minutes before each self-stimulation session. On the other 4 consecutive days, animals received an intraperitoneal injection of sterile saline 30 minutes before the self-stimulation. For both enkephalin-cre and dynorphin-cre cohorts the experiment was counterbalanced, with half of the animals receiving CNO on the first four days after training followed by 4 sessions with sterile saline pretreatment. The other half of the cohorts received intraperitoneal saline during the first four self-stimulation sessions post-training and CNO pretreatment 30 minutes before the last four self-stimulation sessions.

#### REAL-TIME PLACE TEST (RTPT)

All behaviors were performed within a sound-attenuated room maintained at 23°C after habituation to the holding room and the final surgery for at least 1 week. Movements were video recorded and analyzed using AnyMaze software (Stoelting Co). After recovery from both viral and fiber implants surgeries (see above section for the procedure), mice were gently placed in a custom-made unbiased and balanced two-compartment conditioning apparatus (52.5 × 25.5 × 25.5 cm) as described previously ^38,49,61^. For **Figure 1E-H**, the RTPT lasted for 40 minutes, where entry into one compartment triggered light stimulation (20 Hz, 5 ms pulse width, 4mW). Light stimulation was delivered as long as the animal remained in this compartment and would only stop upon entry into the other compartment. The light stimulation paired compartment was also swapped half-way through the trial (20 min after trial starts) to assess animals’ flexibility in seeking light stimulation and confirm its reinforcing value.

### BRAIN CLEARING

Sample Collection: Mice that received a bilateral injection of AAV1.Syn.NES.jRGECO1a.WPRE.SV40 (Penn Vector Core, University of Pennsylvania, viral titer: 2.95 × 10^13^ vg diluted 1:1 with PBS) in dCA1 were deeply anesthetized with isofluorane and transcranially perfused with 4% PFA/PBS. The whole brain was extracted and post-fixed overnight in 4% PFA, then transferred to 30% sucrose in PBS.

Sample Pretreatment with Methanol: Whole brains were cut into halves on the sagittal plane and trimmed to appropriate size. Each half was dehydrated with a series of methanol dilutions in H2O: 20%, 40%, 60%, 80%, and 100%, incubated in 66% dichloromethane/33% methanol, washed with 100% methanol, bleached overnight with chilled 5% H_2_O_2_ in methanol at 4°C, rehydrated with the methanol/H_2_O series: 80%, 60%, 40%, 20%, and washed in PBS/TritonX-100.

Immunolabeling: Brains were treated with a permeabilization solution (PBS, TritonX-100, Glycine, and DMSO), blocked with blocking buffer (6% Donkey Serum in PBS, TritonX-100, and DMSO), then stained with 1:1000 Rabbit Anti-RFP (Rockland Immunochemicals, #600-401-379) primary antibody overnight (5% Donkey Serum in PBS, Tween-20, Heparin, and DMSO) & 1:500 Donkey Anti-Rabbit 647 (Fisher, #A-31573) secondary antibody (3% Donkey Serum in PBS, Tween-20, Heparin).

Clearing: Dehydrate brains with methanol dilutions in H_2_O: 20%, 40%, 60%, 80%, and 100%, continue washing with 100% methanol, then incubate the tissue in the following three solutions consecutively: 66% dichloromethane/33% methanol, 100% dichloromethane, and 100% dibenzyl ether.

Imaging: Cleared brains were imaged at the UNC Microscopy Services Laboratory using the Lavision Ultramicroscope II light-sheet system and visualized with the Imaris software.

### IMAGING

Mice were anesthetized with 1-3% isofluorane and head-fixed in a stereotaxic apparatus (Stoelting). 500 nl of AAV5-CaMKII-ChR2-eYFP (Hope Center Viral Vector Core, viral titer 2.8 × 10^13^ vg/mL) virus was infused bilaterally into the dHPC (rate of infusion: 100 nl/min. Coordinates: A/P: –1.7 mm, M/L: ±1.5 mm, D/V: –1.8 mm). Mice were transcardially perfused with ice-cold 4% paraformaldehyde (PFA) in phosphate-buffered saline (PBS) while deeply anesthetized with isofluorane. The whole brain was extracted and post-fixed overnight in 4% PFA before the brains were transferred for equilibration in 30% sucrose in PBS. Equilibrated brains were flash-frozen and the entire brain was sectioned into 40 μm slices using a cryostat. Floating slices were washed 3×5 min with PBS, blocked with blocking buffer (4% normal donkey serum, 4% bovine serum albumin, and 0.5% TX-100 in PBS) and stained with 1:1,000 Chicken anti-GFP (Abcam #ab13970, RRID: AB_300798) primary antibody overnight. Sections were washed 3×5 min with PBS and stained with 1:1,000 donkey anti-chicken Alexa Fluor 488 secondary antibody (Jackson ImmunoResearch #703-545-155, RRID: AB_2340375), rinsed with PBS, and dried. Slides were cover slipped with hardset antifade mounting media with DAPI (Vectashield) and imaged via confocal microscopy (Leica) or an AxioScan Z1 slide scanning microscope (Zeiss).

### LOCAL FIELD POTENTIAL RECORDINGS

#### CUSTOM-MADE ELECTRODE AND HEAD CAP

The electrode was made in-house using 16 tungsten wires (0.0015”, coated, California Fine Wire) for recording and 1 silver wire (0.010”, coated, California Fine Wire) as ground. The ground wire was soldered on an 18 pin, 16 channel electrical interface board (EIB) (Triangle BioSystems Inc.) while the recording wires (8 on the left and 8 on the right side of the EIB) were connected using small gold EIB pins (Triangle Biosystems Inc.). The electrode was then secured to a 3D-printed head cap that was designed to hold both the electrode and optic fiber at the desired coordinates during implant. Optic fibers (Thorlabs) were cleaved at a length that allow 2.1 mm protruding out from the bottom of the head cap. The 8 tungsten wires on each side are then twisted and pulled to make them straight. The wires were then cut to the desired length (5 mm) before implant.

#### SURGICAL IMPLANT OF THE ELECTRODES

Mice were anesthetized in an induction chamber (4 MAC Isoflurane) and placed into a stereotaxic frame (Kopf Instruments, Model 1900) where they were maintained at 1 MAC–2 MAC isoflurane. Anesthesia was confirmed with a toe pinch. Animals were mounted on a stereotaxic frame in sterile conditions (Stoelting). A total of four holes were drilled into the skull, one above the left dorsal hippocampus (A/P: – 1.70mm, M/L: ± 1.5mm), two above the nucleus accumbens (A/P: + 1.34mm, M/L: ± 0.8mm) and one to secure the ground screw and wire. A unilateral injection, using a blunt needle (86200, Hamilton Company), of AAV5-CaMKII-ChR2-eYFP (500 nl, Hope Center Viral Vector Core, viral titer 2.3 × 10^13^ vg/mL) into dHPC (D/V: –1.80mm; Fakira et al., 2016). After viral injection, the skull was gently scratched across and the custom-made headcap carefully aligned before slowly lowered to the depth desired (1.80 mm DV for dHPC and 4.75 mm DV for NAc). The headcap were secured and affixed to the skull with Metabond dental cement using Radiopaque L-powder, B Quick Base, and C Universal TBB Catalyst (Parkell). Animals were allowed to recover for 7 days prior to conditioning with gentle handling.

#### EXPERIMENT LFP PROTOCOL

Mice were habituated to a black opaque testing chamber (17 cm width x 30.5 cm length x 25 cm height) and the electrophysiology recording setup for 3 consecutive days before test day. 1 day prior to the test day (3^rd^ day of habituation), recording of the evoked local field potential (LFP) was taken to ensure that there was a stable response in the NAc with the activation of the dHPC CaMKII+ neurons. On test day, mice were connected to a head-stage amplifier (Plexon model HST/16D Gen2) and PLEXON amplifier (Plexon Omniplex-D acquisition system) and allowed to habituate for 30 minutes before the start of recording. The neuronal recordings were collected using the OMNIPLEX PLEXON neurophysiology acquisition system and software (Plexon Inc). The dHPC was stimulated for a duration of 5 ms every 5 s while evoked LFP in the NAc (bilateral) was recorded. The peri histogram of evoked LFP recorded in the NAc was analyzed using Neuroexplorer (Plexon Inc).

### FIBER PHOTOMETRY

For fiber photometry studies, an optic fiber was attached to the implanted fiber using a ferrule sleeve (Doric, ZR_2.5). Two LEDs were used to excite GCaMP6f in WT or cre-dependent mice. A 531-Hz sinusoidal LED light (Thorlabs, LED light: M470F3; LED driver: DC4104) was bandpass filtered (470 ± 20 nm, Doric, FMC4) to excite GCaMP6f and evoke Ca2+-dependent emission. A 211-Hz sinusoidal LED light (Thorlabs, LED light: M405FP1; LED driver: DC4104) was bandpass filtered (405 ± 10 nm, Doric, FMC4) to excite GCaMP6f and evoke Ca2+-independent isosbestic control emission. Laser intensity for the 470 nm and 405 nm wavelength bands were measured at the tip of the optic fiber and adjusted to 50 μW before each day of recording. GCaMP6f fluorescence traveled through the same optic fiber before being bandpass filtered (525 ± 25 nm, Doric, FMC4), transduced by a femtowatt silicon photoreceiver (Newport, 2151) and recorded by a real-time processor (TDT, RZ5P). The envelopes of the 531-Hz and 211-Hz signals were extracted in real-time by the TDT program Synapse at a sampling rate of 1017.25 Hz.

Mice received sustained 2 seconds non-contingent stimulation in dCA1 (20 Hz, 5 ms, every 30 seconds) to stimulate ChR2 expressing CaMKII+ neurons within the dorsal hippocampus. The same mice were also trained in operant conditioning (Med Associates Inc.) as described in behavioral paradigm above (light-stimulation). When self-stimulation was consistent across session, fiber photometry recordings were made throughout one-hour operant conditioning sessions. All signals were aligned to light-stimulation delivery, or cue light turning back on after time out period as a control. The resulting signal recorded by TDT Synapse program was analyzed using a custom-written MATLAB scripts. In brief, signals from the isosbestic 405 nm control channel were smoothed and regressed on the smoothed GCaMP 470 nm signal to generate a predicted 405 nm channel using a linear model generated during the regression. Changes in fluorescence across the experiment session (ΔF) were then calculated by subtracting the predicted 405 nm from the raw GCaMP signal to remove photo-bleaching and fiber-bending artifacts. Signals from the GCaMP channel were then divided by the control signal to generate the ΔF/F. Peri-event histograms (Z-score) were then created by averaging changes in fluorescence (ΔF/F) across repeated trials using duration window that encompassed events of interest (stimulation or cue) ^62^.

### QUANTIFICATION AND STATISTICAL ANALYSIS

All experiments were performed at least twice, including all treatment groups in each round, to avoid any unspecific day/condition effect. Treatments (i.e., choice of viral compounds within the dHPC and NAcSh) were randomly assigned to animals before testing. Statistical significance was taken as *p < 0.05, **p < 0.01, ***p < 0.001, and ****p < 0.0001, as determined by a one-way ANOVA or a two-way repeated-measures ANOVA followed by a Sidak post hoc tests, two-tailed unpaired or paired t test, one sample t-test compared to hypothetical value (50%) or Mann-Whitney for unpaired values as appropriate. All data samples were tested for normality of distribution using Shapiro-Wilk test before being assigned to ANOVAs, t-test, or Mann-Whitney analysis. All data are expressed as mean ± SEM. Sample size (n number) always refers to the value obtained from an individual animal when referring to behavioral experiments and anatomical slice analysis. For each experiment detailed statistical analysis and sample size (n number) are carefully reported in supplementary table and figure legends. Statistical analyses were performed in GraphPad Prism 9.0 and SPSS.

### DATA AND SOFTWARE AVAILABILITY

Authors can confirm that all relevant data are included in the paper and its supplementary information files.

## ACKNOWLEDGMENTS

We would like to thank Dr. Alexxai Kravitz for his critical help in experimental design and contribution in acquiring and analyzing local field potential data. In addition, we would like to thank Ms. Ashley Park and all members from the Moron-Concepcion’s lab for their help throughout the completion of the current study. This work was supported by US National Institutes of Health (NIH) grant DA041781 (J.A.M.), DA042499 (J.A.M.), DA041883 (J.A.M.), DA045463 (J.A.M.), DA055047 (N.M.), NARSAD Independent Investigator Award from the Brain and Behavior Research Foundation (J.A.M.), Philippe Foundation (N.M.), McDonnell Center for Cellular and Molecular Neurobiology (N.M.).

## AUTHORS CONTRIBUTIONS

Conceptualization: N.M., K.M.I., T.L.K, J.A.M.; Methodology, N.M., K.M.I. and J.A.M.; Formal analysis: N.M., K.M.I. and J.A.M.; Investigation: N.M., K.M.I., H.J.Y., R.S., S.W., H.F., W.P., W.Y., O.I., A.Z., S.P. ; Writing - Original Draft: N.M., K.M.I. and J.A.M.; Writing – Review & Editing: N.M., K.M.I, H.Y.J., S.W., H.F., T.L.K. and J.A.M.; Funding Acquisition : N.M., T.L.K. and J.A.M.; Resources : T.L.K., and J.A.M.; Supervision: N.M., K.M.I., T.L.K. and J.A.M.

## DECLARATION OF INTERESTS

The authors declare no competing financial interests.

## MAIN FIGURES TITLES AND LEGENDS

**Suppl Figure 1.**
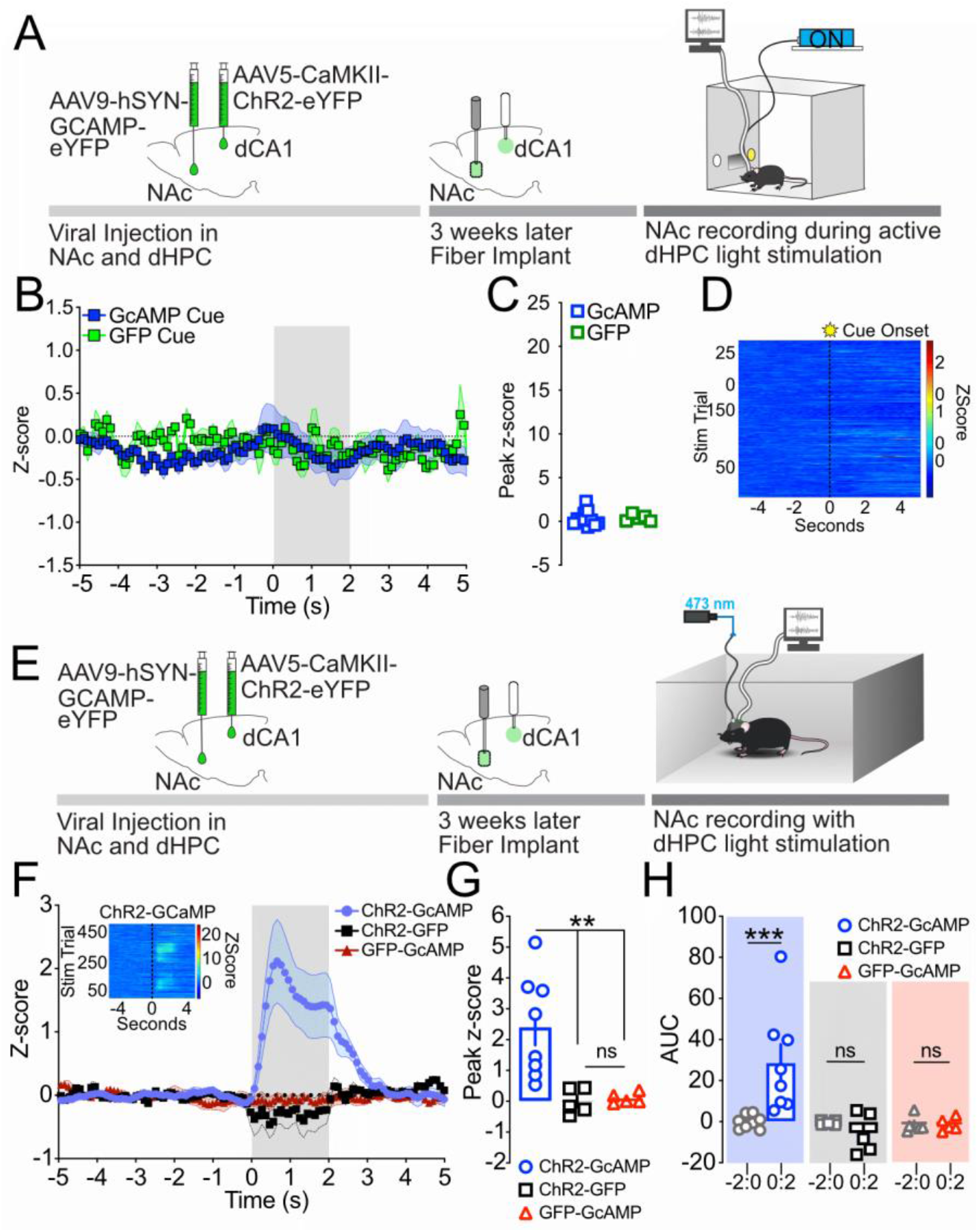
Increase in NAc calcium transients is dependent on dCA1 activation, but not cue associated with light stimulation. A) Representative schematic of the behavior. B) Time course for the Z-scores of the calcium transient. Gray area represents cue light onset. Calcium transients are not impacted by cue light delivery. C) Peak Z-score during the 2 seconds after cue light onset. Peak Z-score remains unchanged in both GCaMP and GFP expressing animals upon cue light onset. D) Heatmap of the calcium transient upon cue light onset. Data are expressed as mean +/-S.E.M. and n.s. p > 0.05. T-test. E) Schematic representation of the experimenter-induced stimulation procedure. F) Time course for the Z-scores of the calcium transient. Gray area represents dCA1 stimulation. Calcium transients selectively increase in animals expressing both ChR2 in the dCA1 and GCaMP6f in the NAcSh upon stimulation of CaMKII+ neurons. Insert: heatmap of the calcium transient upon light delivery. G) Peak Z-score during the 2 seconds of dCA1 CaMKII+ neurons stimulation. Peak Z-score is significantly higher in ChR2-GCaMP6f expressing animals. H) Area under the curve (AUC) of 2 seconds prior and during dCA1 light stimulation obtained in experimental groups. The AUC of the calcium transients measured in the NAc of ChR2-GCaMP animals were significantly higher during stimulation (ChR2-GCaMP: 0:2) compared to their respective baseline (ChR2-GCaMP: -2:0). The AUC of the calcium transients recorded did not significantly change upon light stimulation in the dCA1 in negative control groups (ChR2-GFP and GFP-GCaMP). Data are expressed as mean +/-S.E.M. and n.s. p > 0.05; ** p < 0.001; and *** p < 0.001

**Suppl Figure 2:**
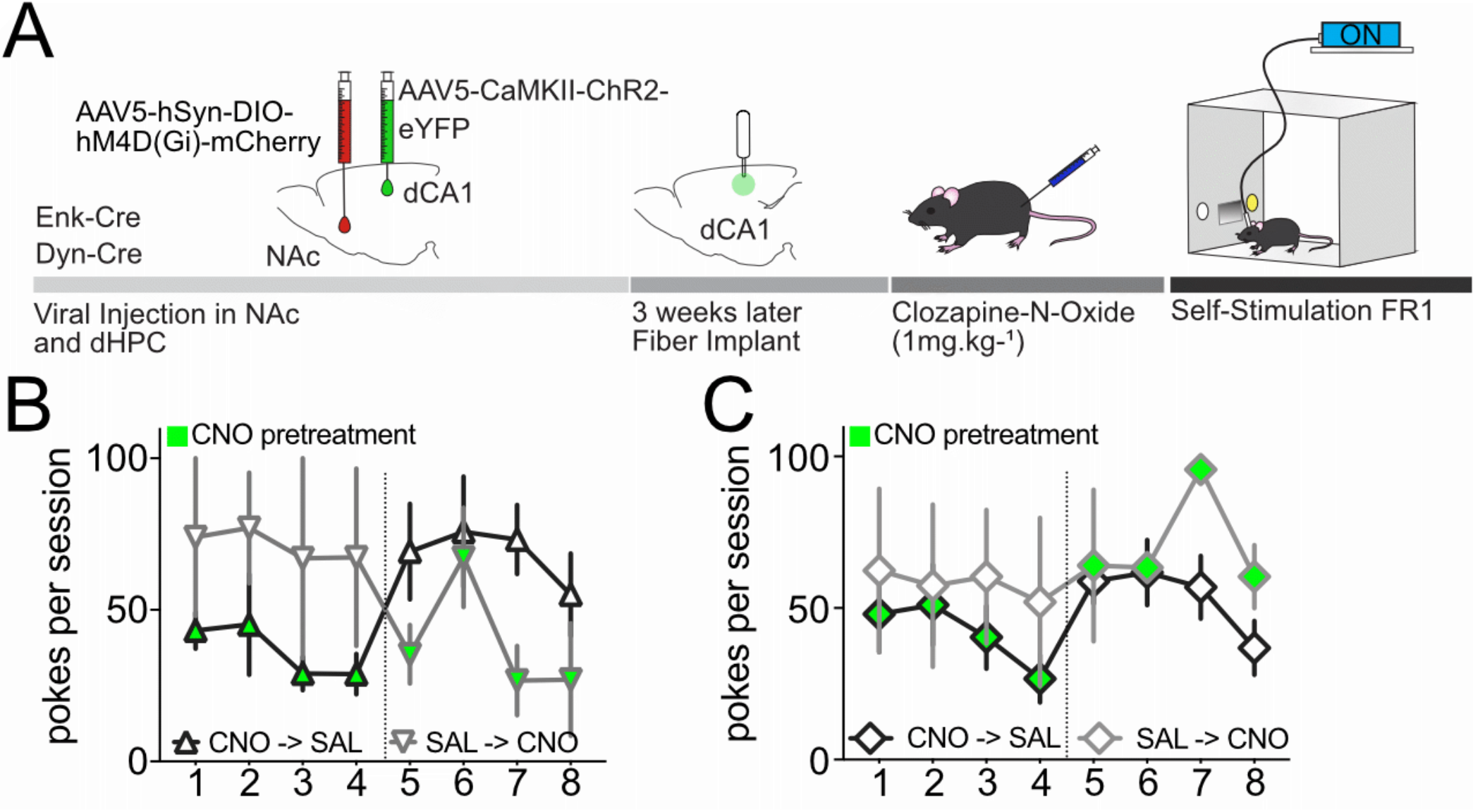
Silencing dynorphin containing neurons, but not enkephalin, decreases active light self-stimulation. A) Representative schematic of the behavior. B) Daily representation of nose pokes in the active inset per session for Dyn-cre animals. While one group of animals received 4 days of saline i.p. pretreatment before being exposed to 4 sessions with CNO i.p. pretreatment (grey downwards triangles), the other group was first exposed to four days of CNO before receiving saline i.p. pretreatment (black upwards triangles). Regardless of the days animals received CNO pre-treatment, silencing dynorphin containing neurons reduces the number of pokes in the active inset (self-stimulation). C) Daily representation of nose pokes in the active inset per session for Enk-cre animals. Animals were exposed to a similar saline/CNO pretreatment schedule as mentioned above. However, silencing NAc enkephalin-containing does not impact nose pokes in the active inset (self-stimulation). Data are expressed as mean +/-S.E.M.

**Suppl Figure 3:**
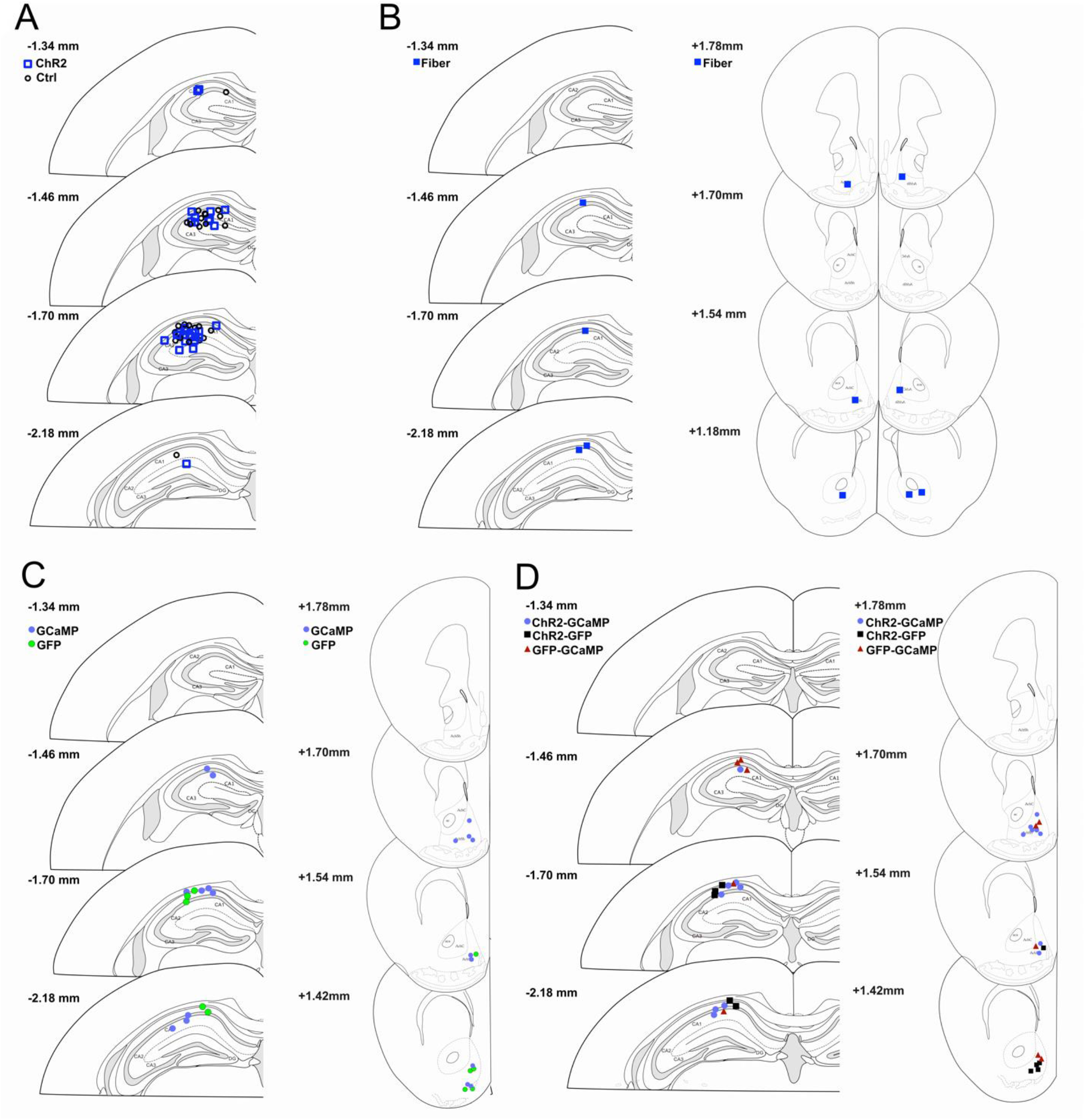
Histology Figures 1 and 2. A) Histology for all fiber implants described in Figure 1. B) Histology for fiber implants located in the dCA1 and the recording electrodes in the NAc for Figure 2 D-E. C) Histology for Fiber implants in the dCA1 (Stim) and the NAc (recording) for Figure 2 H-J and Suppl Figure 1 B-D. D) Histology for Fiber implants in the dCA1 (Stim) and the NAc (recording) for Suppl Figure 1 E-H.

**Suppl Figure 4:**
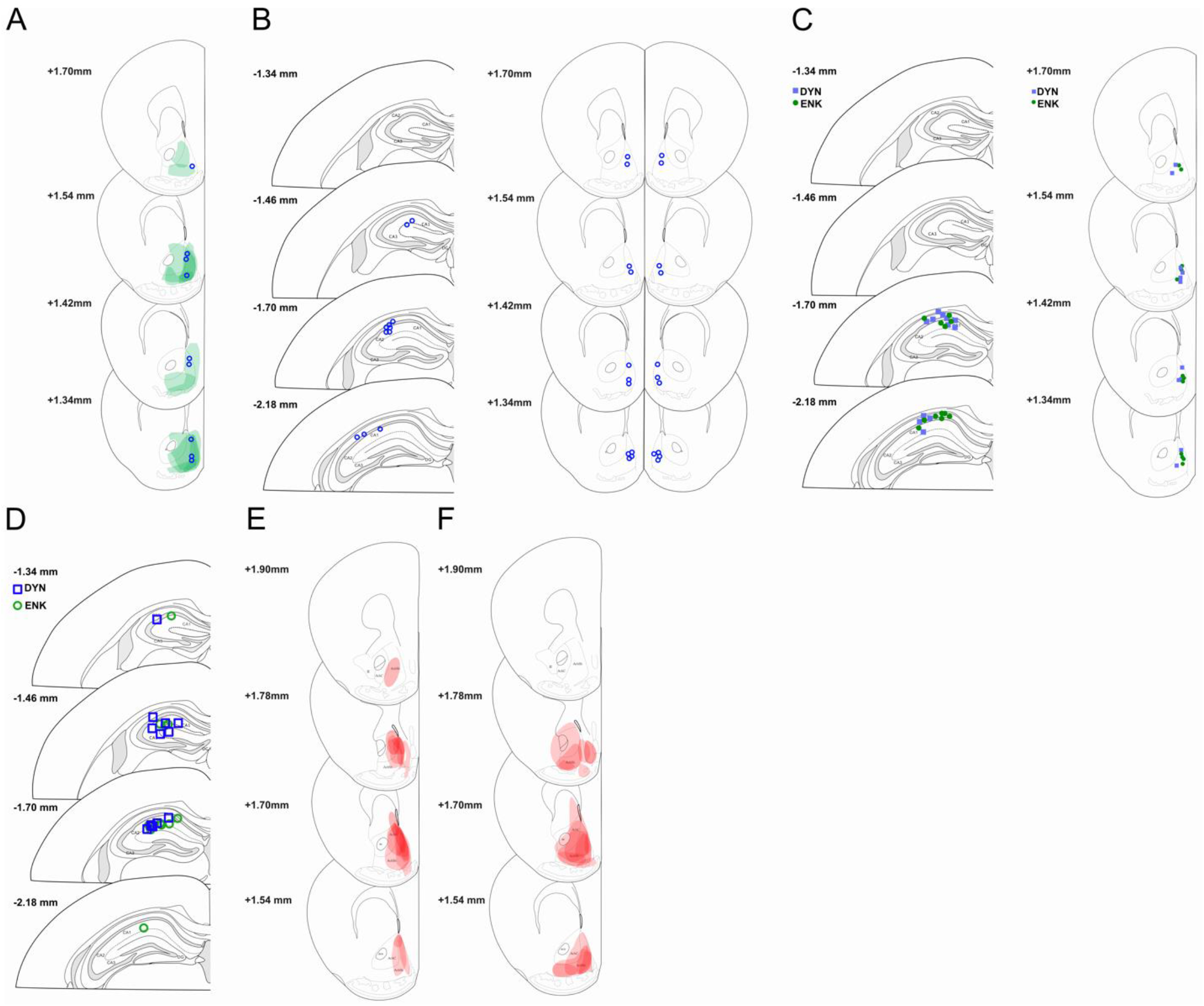
Histology Figures 3 and 4. A) Histology for all fiber implants and ChR2 expression in the NAc described in Figure 3 B-C. B) Histology for injection sites of AP5/CNQX in the NAc for Figure 3 E-F. C) Histology for Fiber implants in the dCA1 (Stim) and the NAc (recording) for Figure 4 B-D. D) Histology for Fiber implants in the dCA1 (Stim) Figures 4 F-H. E) Histology for Gi DREAAD expression in Dyn+ neurons in the NAc for Figure 4 F. F) Histology for Gi DREAAD expression in Enk+ neurons in the NAc for Figure 4 F.

## Notes

### Competing Interest Statement

The authors have declared no competing interest.

## REFERENCES

1. Moser, M. B., Moser, E. I., Forrest, E., Andersen, P. & Morris, R. G. Spatial learning with a minislab in the dorsal hippocampus. Proc Natl Acad Sci U S A 92, 9697–9701 (1995).

2. Strange, B. A., Witter, M. P., Lein, E. S. & Moser, E. I. Functional organization of the hippocampal longitudinal axis. Nat Rev Neurosci 15, 655–669 (2014).

3. Yang, A. K., Mendoza, J. A., Lafferty, C. K., Lacroix, F. & Britt, J. P. Hippocampal Input to the Nucleus Accumbens Shell Enhances Food Palatability. Biol Psychiatry 87, 597–608 (2020).

4. Yoshida, K., Drew, M. R., Mimura, M. & Tanaka, K. F. Serotonin-mediated inhibition of ventral hippocampus is required for sustained goal-directed behavior. Nat Neurosci 22, 770–777 (2019).

5. Yoshida, K. et al. Chronic social defeat stress impairs goal-directed behavior through dysregulation of ventral hippocampal activity in male mice. Neuropsychopharmacology 46, 1606–1616 (2021).

6. Adhikari, A., Topiwala, M. A. & Gordon, J. A. Synchronized Activity between the Ventral Hippocampus and the Medial Prefrontal Cortex during Anxiety. Neuron 65, 257–269 (2010).

7. Jimenez, J. C. et al. Anxiety Cells in a Hippocampal-Hypothalamic Circuit. Neuron 97, 670-683.e6 (2018).

8. Lasseter, H. C., Xie, X., Ramirez, D. R. & Fuchs, R. A. Sub-region specific contribution of the ventral hippocampus to drug context-induced reinstatement of cocaine-seeking behavior in rats. Neuroscience 171, 830–839 (2010).

9. LeGates, T. A. et al. Reward behavior is regulated by the strength of hippocampus-nucleus accumbens synapses. Nature 564, 258–262 (2018).

10. Marek, R. et al. Hippocampus-driven feed-forward inhibition of the prefrontal cortex mediates relapse of extinguished fear. Nat Neurosci 21, 384–392 (2018).

11. Pignatelli, M. et al. Cooperative synaptic and intrinsic plasticity in a disynaptic limbic circuit drive stress-induced anhedonia and passive coping in mice. Molecular Psychiatry 1–20 (2020) doi:10.1038/s41380-020-0686-8.

12. Sweeney, P. & Yang, Y. An excitatory ventral hippocampus to lateral septum circuit that suppresses feeding. Nat Commun 6, 10188 (2015).

13. Wang, N. et al. Role of Glutamatergic Projections from the Ventral CA1 to Infralimbic Cortex in Context-Induced Reinstatement of Heroin Seeking. Neuropsychopharmacology 43, 1373–1384 (2018).

14. Komorowski, R. W. et al. Ventral hippocampal neurons are shaped by experience to represent behaviorally relevant contexts. J Neurosci 33, 8079–8087 (2013).

15. Moser, M. B. & Moser, E. I. Functional differentiation in the hippocampus. Hippocampus 8, 608–619 (1998).

16. Billa, S. K., Sinha, N., Rudrabhatla, S. R. & Morón, J. A. Extinction of morphine-dependent conditioned behavior is associated with increased phosphorylation of the GluR1 subunit of AMPA receptors at hippocampal synapses. Eur J Neurosci 29, 55–64 (2009).

17. Billa, S. K., Xia, Y. & Morón, J. A. Disruption of morphine conditioned place preference by delta-2-opioid receptor antagonist: Study of mu- and delta-opioid receptor expression at the synapse. Eur J Neurosci 32, 625–631 (2010).

18. Chersi, F. & Burgess, N. The Cognitive Architecture of Spatial Navigation: Hippocampal and Striatal Contributions. Neuron 88, 64–77 (2015).

19. Fakira, A. K., Portugal, G. S., Carusillo, B., Melyan, Z. & Morón, J. A. Increased small conductance calcium-activated potassium type 2 channel-mediated negative feedback on N-methyl-D-aspartate receptors impairs synaptic plasticity following context-dependent sensitization to morphine. Biol Psychiatry 75, 105–114 (2014).

20. Ito, R., Robbins, T. W., Pennartz, C. M. & Everitt, B. J. Functional interaction between the hippocampus and nucleus accumbens shell is necessary for the acquisition of appetitive spatial context conditioning. J Neurosci 28, 6950–6959 (2008).

21. Portugal, G. S. et al. Hippocampal Long-Term Potentiation Is Disrupted during Expression and Extinction But Is Restored after Reinstatement of Morphine Place Preference. J Neurosci 34, 527–538 (2014).

22. Trouche, S. et al. Recoding a cocaine-place memory engram to a neutral engram in the hippocampus. Nat Neurosci 19, 564–567 (2016).

23. Trouche, S. et al. A Hippocampus-Accumbens Tripartite Neuronal Motif Guides Appetitive Memory in Space. Cell 176, 1393-1406.e16 (2019).

24. Xia, L., Nygard, S. K., Sobczak, G. G., Hourguettes, N. J. & Bruchas, M. R. Dorsal-CA1 Hippocampal Neuronal Ensembles Encode Nicotine-Reward Contextual Associations. Cell Rep 19, 2143–2156 (2017).

25. Xia, Y. et al. Hippocampal GluA1-Containing AMPA Receptors Mediate Context-Dependent Sensitization to Morphine. J. Neurosci. 31, 16279–16291 (2011).

26. Campese, V. D. & Delamater, A. R. Dorsal hippocampus inactivation impairs spontaneous recovery of Pavlovian magazine approach responding in rats. Behav Brain Res 269, 37–43 (2014).

27. O’Keefe, J. & Dostrovsky, J. The hippocampus as a spatial map. Preliminary evidence from unit activity in the freely-moving rat. Brain Res 34, 171–175 (1971).

28. Ramirez, D. R. et al. Dorsal Hippocampal Regulation of Memory Reconsolidation Processes that Facilitate Drug Context-induced Cocaine-seeking Behavior in Rats. Eur J Neurosci 30, 901–912 (2009).

29. Ye, X., Kapeller-Libermann, D., Travaglia, A., Inda, M. C. & Alberini, C. M. Direct dorsal hippocampal-prelimbic cortex connections strengthen fear memories. Nat Neurosci 20, 52–61 (2017).

30. Berridge, K. C. & Robinson, T. E. What is the role of dopamine in reward: hedonic impact, reward learning, or incentive salience? Brain Res. Brain Res. Rev. 28, 309–369 (1998).

31. Everitt, B. J. & Robbins, T. W. Neural systems of reinforcement for drug addiction: from actions to habits to compulsion. Nat Neurosci 8, 1481–1489 (2005).

32. Wise, R. A. Dopamine, learning and motivation. Nat Rev Neurosci 5, 483–494 (2004).

33. Britt, J. P. et al. Synaptic and behavioral profile of multiple glutamatergic inputs to the nucleus accumbens. Neuron 76, 790–803 (2012).

34. Ursin, R., Ursin, H. & Olds, J. Self-stimulation of hippocampus in rats. J Comp Physiol Psychol 61, 353–359 (1966).

35. Van Der Kooy, D., Fibiger, H. C. & Phillips, A. G. Monoamine involvement in hippocampal self-stimulation. Brain Research 136, 119–130 (1977).

36. Mahler, S. V., Smith, R. J. & Aston-Jones, G. Interactions between VTA orexin and glutamate in cue-induced reinstatement of cocaine seeking in rats. Psychopharmacology (Berl) 226, 687–698 (2013).

37. Castro, D. C. & Bruchas, M. R. A Motivational and Neuropeptidergic Hub: Anatomical and Functional Diversity within Nucleus Accumbens Shell. Neuron 102, 529–552 (2019).

38. Massaly, N. et al. Pain-Induced Negative Affect Is Mediated via Recruitment of The Nucleus Accumbens Kappa Opioid System. Neuron 102, 564-573.e6 (2019).

39. Alme, C. B. et al. Place cells in the hippocampus: Eleven maps for eleven rooms. PNAS 111, 18428–18435 (2014).

40. Dragoi, G., Harris, K. D. & Buzsáki, G. Place Representation within Hippocampal Networks Is Modified by Long-Term Potentiation. Neuron 39, 843–853 (2003).

41. Robinson, N. T. M. et al. Targeted Activation of Hippocampal Place Cells Drives Memory-Guided Spatial Behavior. Cell 183, 1586-1599.e10 (2020).

42. Lansink, C. S. et al. Reward Expectancy Strengthens CA1 Theta and Beta Band Synchronization and Hippocampal-Ventral Striatal Coupling. J Neurosci 36, 10598–10610 (2016).

43. van der Meer, M. A. A., Johnson, A., Schmitzer-Torbert, N. C. & Redish, A. D. Triple dissociation of information processing in dorsal striatum, ventral striatum, and hippocampus on a learned spatial decision task. Neuron 67, 25–32 (2010).

44. Sjulson, L., Peyrache, A., Cumpelik, A., Cassataro, D. & Buzsáki, G. Cocaine Place Conditioning Strengthens Location-Specific Hippocampal Coupling to the Nucleus Accumbens. Neuron 98, 926-934.e5 (2018).

45. Sosa, M., Joo, H. R. & Frank, L. M. Dorsal and Ventral Hippocampal Sharp-Wave Ripples Activate Distinct Nucleus Accumbens Networks. Neuron 105, 725-741.e8 (2020).

46. O’Donnell, P. & Grace, A. A. Synaptic interactions among excitatory afferents to nucleus accumbens neurons: hippocampal gating of prefrontal cortical input. J Neurosci 15, 3622–3639 (1995).

47. Sesack, S. R. & Grace, A. A. Cortico-Basal Ganglia reward network: microcircuitry. Neuropsychopharmacology 35, 27–47 (2010).

48. Stuber, G. D., Britt, J. P. & Bonci, A. Optogenetic modulation of neural circuits that underlie reward seeking. Biol Psychiatry 71, 1061–1067 (2012).

49. Al-Hasani, R. et al. Distinct Subpopulations of Nucleus Accumbens Dynorphin Neurons Drive Aversion and Reward. Neuron 87, 1063–1077 (2015).

50. Calipari, E. S. et al. In vivo imaging identifies temporal signature of D1 and D2 medium spiny neurons in cocaine reward. Proc Natl Acad Sci U S A 113, 2726–2731 (2016).

51. Creed, M. et al. Cocaine Exposure Enhances the Activity of Ventral Tegmental Area Dopamine Neurons via Calcium-Impermeable NMDARs. J Neurosci 36, 10759–10768 (2016).

52. Kravitz, A. V., Tye, L. D. & Kreitzer, A. C. Distinct roles for direct and indirect pathway striatal neurons in reinforcement. Nat Neurosci 15, 816–818 (2012).

53. Lobo, M. K. et al. Cell type-specific loss of BDNF signaling mimics optogenetic control of cocaine reward. Science 330, 385–390 (2010).

54. Pascoli, V. et al. Contrasting forms of cocaine-evoked plasticity control components of relapse. Nature 509, 459–464 (2014).

55. Ren, W. et al. The indirect pathway of the nucleus accumbens shell amplifies neuropathic pain. Nat. Neurosci. 19, 220–222 (2016).

56. Schwartz, N. et al. Chronic pain. Decreased motivation during chronic pain requires long-term depression in the nucleus accumbens. Science 345, 535–542 (2014).

57. MacAskill, A. F., Cassel, J. M. & Carter, A. G. Cocaine exposure reorganizes cell type- and input-specific connectivity in the nucleus accumbens. Nat Neurosci 17, 1198–1207 (2014).

58. Castro, D. C. & Berridge, K. C. Opioid hedonic hotspot in nucleus accumbens shell: mu, delta, and kappa maps for enhancement of sweetness ‘liking’ and ‘wanting’. J. Neurosci. 34, 4239–4250 (2014).

59. Sosa, M. & Giocomo, L. M. Navigating for reward. Nat Rev Neurosci 22, 472–487 (2021).

60. Fakira, A. K., Massaly, N., Cohensedgh, O., Berman, A. & Morón, J. A. Morphine-Associated Contextual Cues Induce Structural Plasticity in Hippocampal CA1 Pyramidal Neurons. Neuropsychopharmacology 41, 2668–2678 (2016).

61. McCall, J. G. et al. CRH Engagement of the Locus Coeruleus Noradrenergic System Mediates Stress-Induced Anxiety. Neuron 87, 605–620 (2015).

62. Barker, D. J. et al. Lateral Preoptic Control of the Lateral Habenula through Convergent Glutamate and GABA Transmission. Cell Rep 21, 1757–1769 (2017).

