## Supplementary Figures 1-4 for "Dorsal Hippocampus To Nucleus Accumbens Projections Drive Reinforcement Via Activation of Accumbal Dynorphin Neurons"

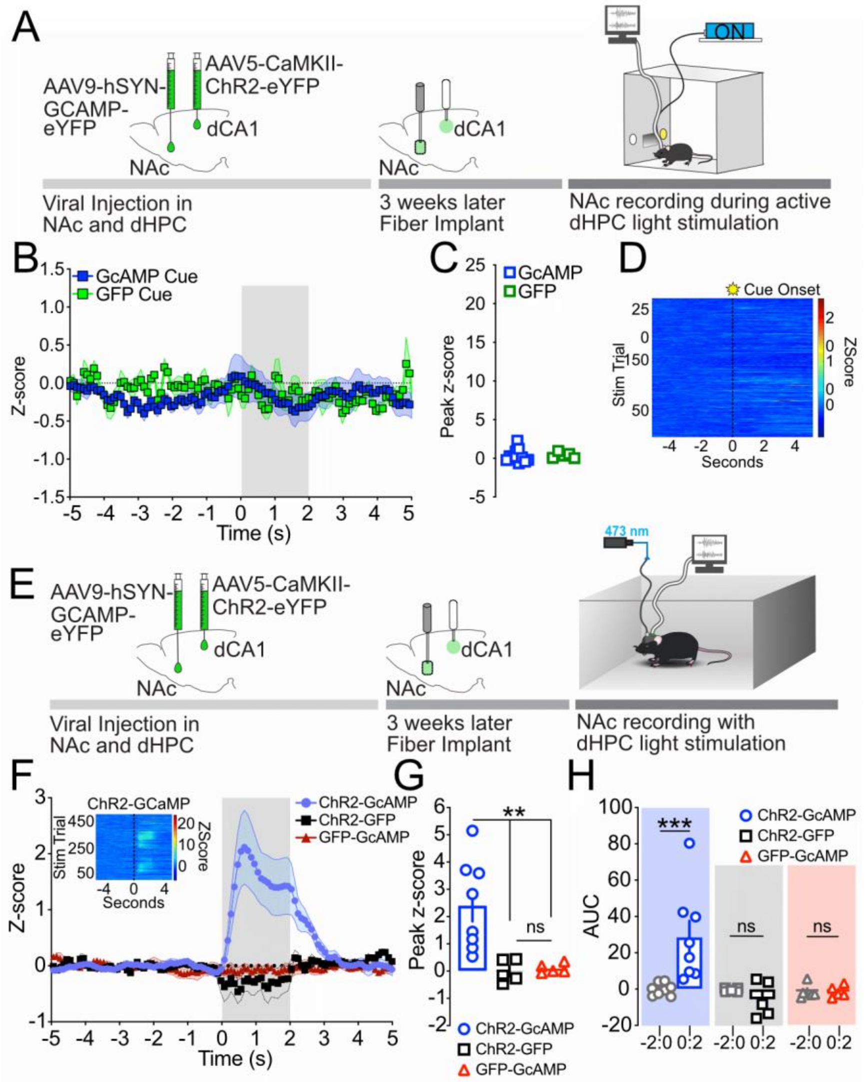


**Suppl Figure 1. Increase in NAc calcium transients is dependent on dCA1 activation, but not cue associated with light stimulation**

A) Representative schematic of the behavior. B) Time course for the Z-scores of the calcium transient. Gray area represents cue light onset. Calcium transients are not impacted by cue light delivery. C) Peak Z-score during the 2 seconds after cue light onset. Peak Z-score remains unchanged in both GCaMP and GFP expressing animals upon cue light onset. D) Heatmap of the calcium transient upon cue light onset. Data are expressed as mean +/- S.E.M. and n.s. $p > 0.05$. T-test. E) Schematic representation of the experimenter-induced stimulation procedure. F) Time course for the Z-scores of the calcium transient. Gray area represents dCA1 stimulation. Calcium transients selectively increase in animals expressing both ChR2 in the dCA1 and GCaMP6f in the NAcSh upon stimulation of CaMKII+ neurons. Insert: heatmap of the calcium transient upon light delivery. G) Peak Z-score during the 2 seconds of dCA1 CaMKII+ neurons stimulation. Peak Z-score is significantly higher in ChR2-GCaMP6f expressing animals. H) Area under the curve (AUC) of 2 seconds prior and during dCA1 light stimulation obtained in experimental groups. The AUC of the calcium transients measured in the NAc of ChR2-GCaMP animals were significantly higher during stimulation (ChR2-GCaMP: 0:2) compared to their respective baseline (ChR2-GCaMP: -2:0). The AUC of the calcium transients recorded did not significantly change upon light stimulation in the dCA1 in negative control groups (ChR2-GFP and GFP-GCaMP). Data are expressed as mean +/- S.E.M. and n.s. $p > 0.05$; ** $p < 0.001$; and *** $p < 0.001$

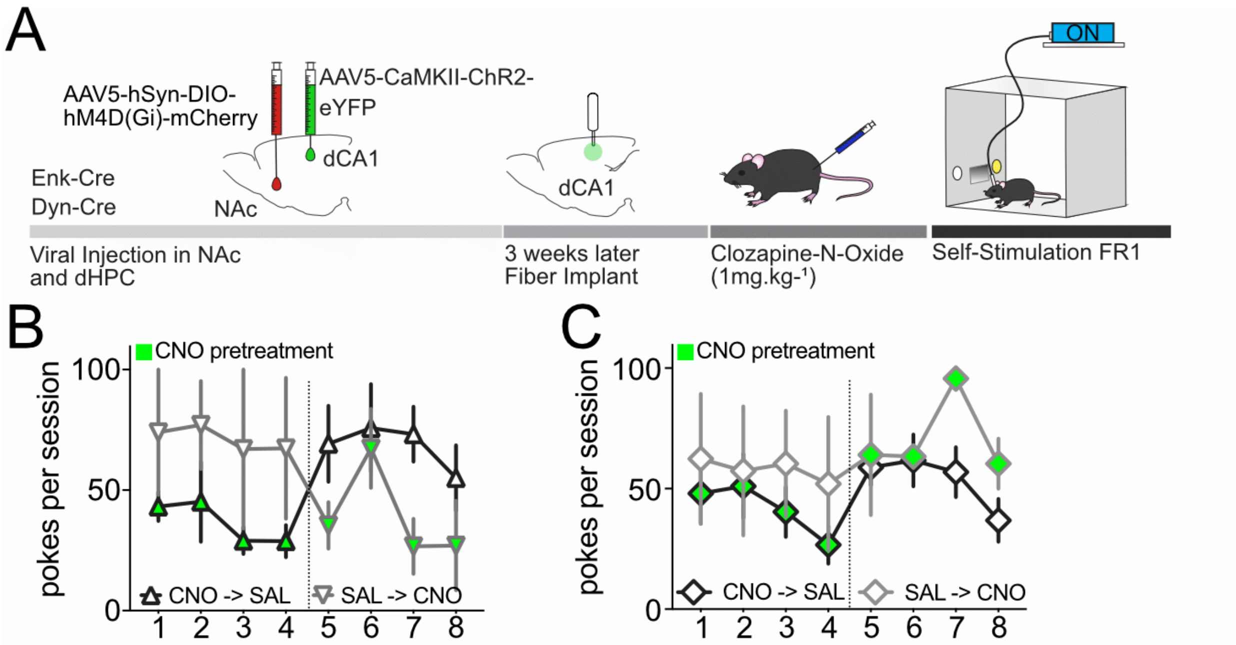


**Suppl Figure 2: Silencing dynorphin containing neurons, but not enkephalin, decreases active light self-stimulation**

A) Representative schematic of the behavior. B) Daily representation of nose pokes in the active inset per session for Dyn-cre animals. While one group of animals received 4 days of saline i.p. pretreatment before being exposed to 4 sessions with CNO i.p. pretreatment (grey downwards triangles) , the other group was first exposed to four days of CNO before receiving saline i.p. pretreatment (black upwards triangles). Regardless of the days animals received CNO pre-treatment, silencing dynorphin containing neurons reduces the number of pokes in the active inset (self-stimulation). C) Daily representation of nose pokes in the active inset per session for Enk-cre animals. Animals were exposed to a similar saline/CNO pretreatment schedule as mentioned above. However, silencing NAc enkephalin-containing does not impact nose pokes in the active inset (self-stimulation). Data are expressed as mean +/- S.E.M.

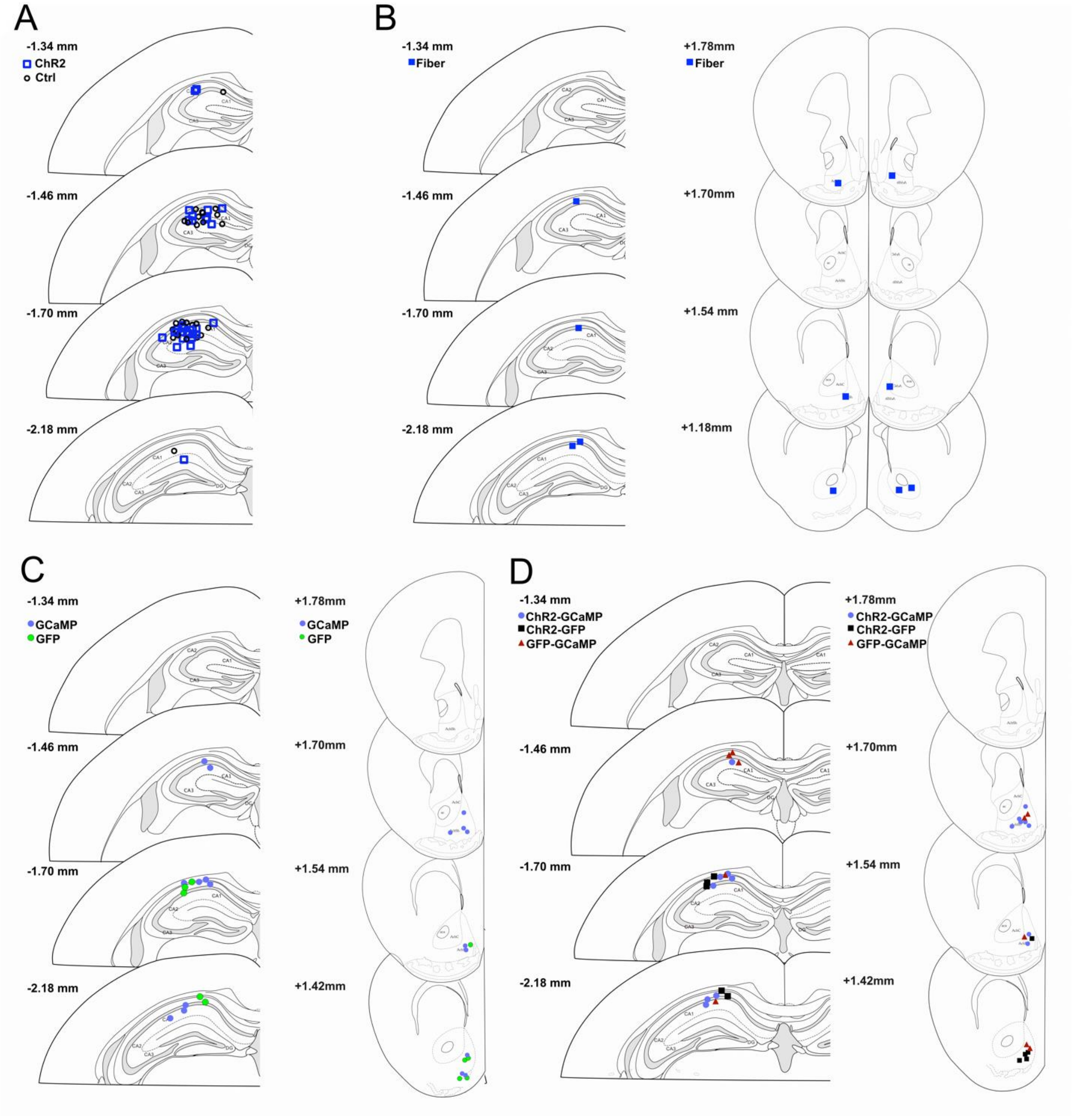


**Suppl Figure 3: Histology Figures 1 and 2**

A) Histology for all fiber implants described in Figure 1. B) Histology for fiber implants located in the dCA1 and the recording electrodes in the NAc for Figure 2 D-E. C) Histology for Fiber implants in the dCA1 (Stim) and the NAc (recording) for Figure 2 H-J and Suppl Figure 1 B-D. D) Histology for Fiber implants in the dCA1 (Stim) and the NAc (recording) for Suppl Figure 1 E-H.

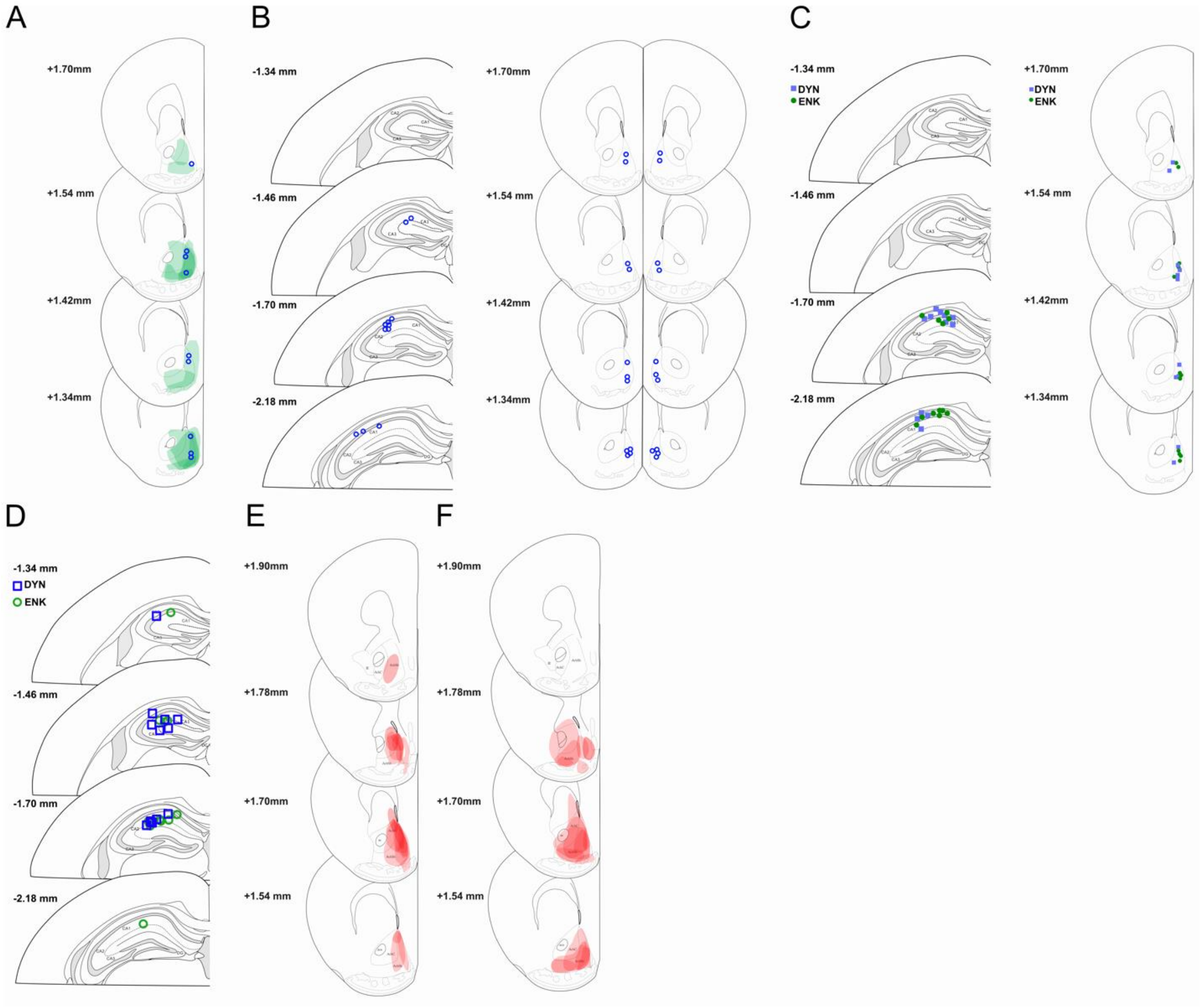


**Suppl Figure 4: Histology Figures 3 and 4**

A) Histology for all fiber implants and ChR2 expression in the NAc described in Figure 3 B-C. B) Histology for injection sites of AP5/CNQX in the NAc for Figure 3 E-F. C) Histology for Fiber implants in the dCA1 (Stim) and the NAc (recording) for Figure 4 B-D. D) Histology for Fiber implants in the dCA1 (Stim) Figures 4 F-H. E) Histology for Gi DREAAD expression in Dyn+ neurons in the NAc for Figure 4 F. F)  Histology for Gi DREAAD expression in Enk+ neurons in the NAc for Figure 4 F.
